# Social foragers rely more on dynamic cues than prior beliefs in unpredictable natural environments

**DOI:** 10.64898/2026.10.30.763071

**Authors:** Alexander Schakowski, Dominik Deffner, Raine Kortet, Petri T. Niemelä, Marwa M. Kavelaars, Maria Pykälä, Ralf H.J.M. Kurvers

## Abstract

Foraging is a prime example of adaptive decision-making in uncertain and changing environments. Foragers need to continuously integrate their direct experiences and observations of what others are doing with what they already know about the distribution of prey. Yet little is known about the importance of prior knowledge relative to dynamically sampled information in social foraging settings. Here, we investigated the role of prior beliefs in five large-scale ice-fishing competitions involving 49–53 highly experienced ice-fishers in 251 foraging trips. To quantify their prior spatial beliefs, participants marked areas they believed to have high resource abundance on topographic maps containing isoclines (i.e., depth curves) before each competition. GPS devices and headcams tracked behavioural states and foraging success. We found that prior beliefs about the fish distribution are to a large extent driven by lake topography and therefore converge across individuals. However, these prior beliefs were not systematically linked to foraging success. While prior beliefs predicted spot choices, especially in the beginning of competitions, they were heavily outweighed by dynamic personal and social information sources. Exploring how the use of prior beliefs differed between foraging trips, we found that high reliance on prior beliefs was associated with reduced avoidance of unsuccessful locations which resulted in limited spatial exploration. We conclude that dynamic information sampling and accumulation are key processes in social foraging decisions, with prior spatial beliefs playing only a minor role in natural environments where resources are hard to predict in advance.

## 1 Introduction

Human cognition has evolved in uncertain and dynamic environments that require individuals to constantly gather and integrate various sources of information to make adaptive decisions [1, 2]. Social foraging, that is, the ability to locate and extract resources in the presence of others, is a prime example of such dynamic decision-making [3–5]. Foragers typically have access to three distinct types of information: (i) personal sampling information (i.e., information obtained through sampling the environment, such as one’s own foraging success), (ii) social information (i.e., information about resource locations provided by the social environment, such as the presence of others) and (iii) prior beliefs or knowledge about the resource environment [2]. Social foraging in particular, thus offers a unique testbed to understand the mechanisms guiding information updating and integration in an ecologically relevant context [3, 4, 6–9].

A rich body of literature across both laboratory and field settings has investigated how human foragers use personal and social information, and how these information sources are integrated during foraging [4, 6, 7, 10–15]. For example, [10] showed that Nahua mushroom foragers use encounter conditional search strategies [11, 16, 17] and decrease their search radius immediately after encountering resources, and [15] provides evidence that humans use social location cues to infer resource quality in a laboratory experiment. [4] and [7] provided evidence that personal and social information are not used independently of each other, but that human foragers use social information conditionally (i.e., using social information only when personal information is unreliable) in both an immersive reality foraging task and under natural conditions in the field. These and other studies have demonstrated the importance of information sampling and of individual and social learning processes for foraging decision-making [4, 6, 18–21].

Particularly in real-world settings information that is sampled during foraging are not the only sources of information available to the decision-maker. Instead, foragers often possess vast knowledge about the foraging ecology, extensive personal experience and complex mental models that guide their information acquisition. Life-history theory, for instance, suggests that a prolonged childhood evolved in humans to allow individuals to gather such complex ecological knowledge and acquire foraging and hunting skills [22–24]. Indeed, research with hunter-gatherer and subsistence horticulturalist societies shows that gathering knowledge (e.g., on the types of prey that habitate the foraging ecology) and foraging skills, for example through cultural transmission [25], is an essential learning goal throughout childhood and an important predictor of foraging success [23, 24, 26].

This raises the question of whether foraging decisions under natural conditions can be better explained by dynamic information sampling or by individuals following strong prior beliefs. To illustrate with an example from non-human animals, elephants’ patch-leaving decisions are best approximated by a Bayesian updating learning model in an experimental setting. In a field setting however, they are best explained by a model assuming perfect knowledge about ecological parameters (e.g., average return rates) [27], suggesting that strong priors might override more recent information under natural conditions.

How foragers integrate prior beliefs about the resource distribution with sampled information to make foraging decisions *in situ* is not well understood, however. Not accounting for such prior beliefs in real-world settings might exaggerate the role of information sampling in foraging decision-making. Building on previous studies that investigated how foraging success, social information and landscape features affected the decision where to search for resources (i.e., spot-selection) and when to leave a location (i.e., spot-leaving decisions) in large-scale ice-fishing competitions in Finland [7, 12, 13], the present study aims to characterize prior spatial beliefs about the resource distribution and how they might relate to mental models about prey behaviour. In addition, we use computational modeling in combination high-resolution GPS tracking and video recordings of large groups of foragers to investigate the extent to which prior beliefs, in contrast to social and personal information, influence where individuals search for resources and when they decide to leave. Ice-fishing competitions—involving mobile and patchy resources, harsh ecological conditions, a large spatial scale, and a rich social and cultural context—offer a naturalistic paradigm to investigate these questions outside the confines of the laboratory.

## 2 Methods

### 2.1 Participants and Data Collection

We briefly summarize the setup of the ice-fishing competitions here and refer to [7] for the full description. During the competitions, large groups of participants compete for three hours to maximize their catch of Eurasian perch (*Perca fluviatilis*). They are free to relocate, drill and fish in a large designated competition area (median = 3.06km^2^, min = 1.18km^2^, max = 3.56km^2^; hereafter we refer to competition areas as lakes) (see also Fig. 1A). After each competition, each participant’s catch is weighed, and they are incentivized based on their ranking (receiving up to

**Figure 1.**
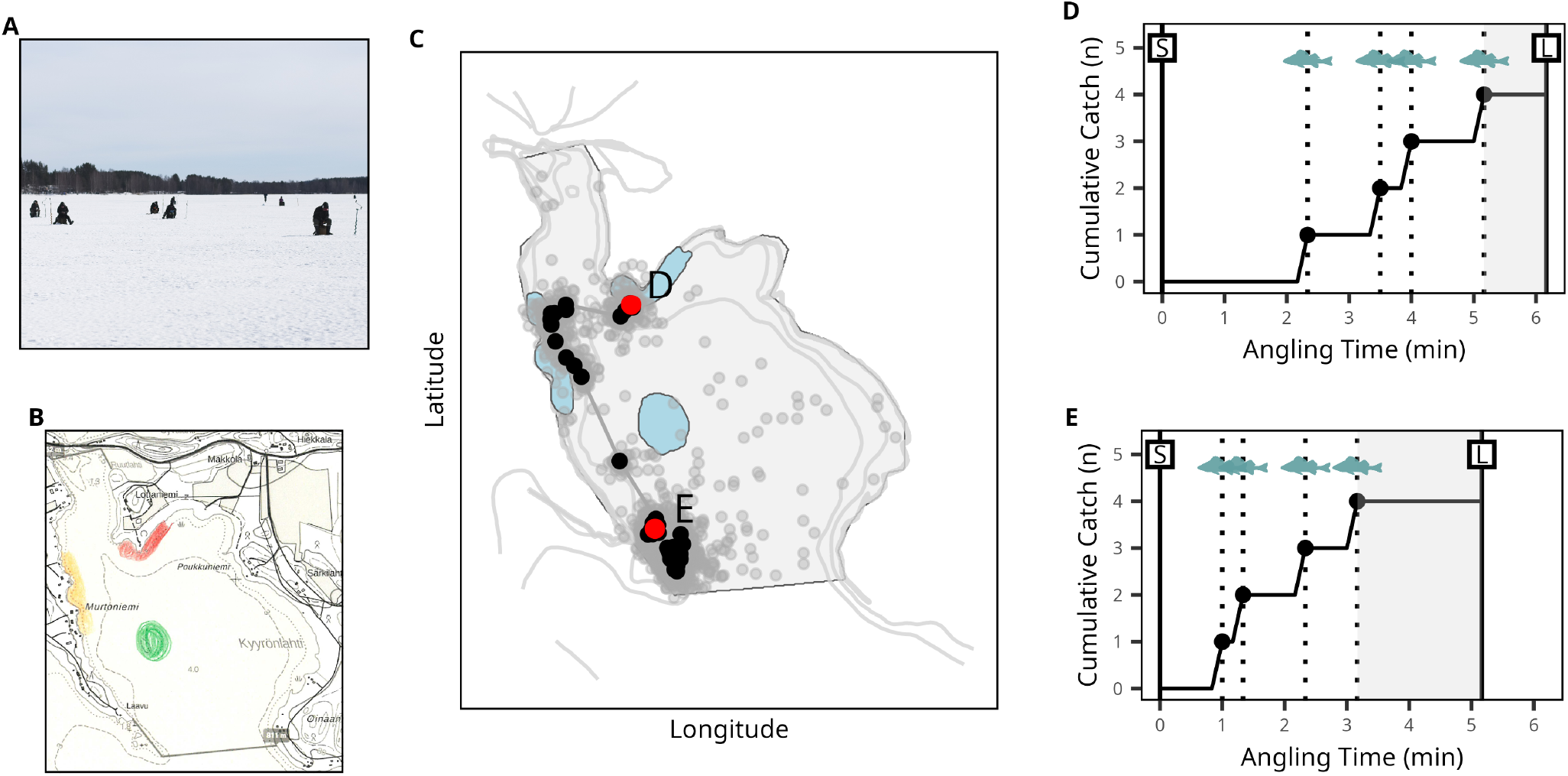
Illustration of the ice-fishing study system, and the methods used to quantify and study the role of prior spatial beliefs in foraging behaviour. (**A**) Aerial image taken during a competition showing a subset of participants relocating, drilling or fishing. (**B**) Before each foraging trip, participants marked a map containing isoclines (contour lines indicating equal depth) with areas they would definitely (red), very likely (yellow), and likely (green) visit when fishing alone to maximize their catch. (**C**) To study whether these prior beliefs affected participants’ spot selection, we compared the distance between the actual angling spots (black) and the nominated areas (light blue) to the distance between simulated spots (grey) and the nominated areas for each spot choice (see Methods). (**D, E**) To study whether prior beliefs affected participants’ spot-leaving decisions, we quantified at each angling spot the start time of the fishing event (S), the leaving time (L), and all fish catches (fish icons). We then estimated whether giving-up times (i.e., the time between catching the last fish and leaving a spot; shaded area) differed between spots closer to nominated areas (D) and further away from nominated areas (E), controlling for catching success.

€500). All participants were equipped with GPS devices logging spatial position every second and headcams continuously recording foraging success (i.e., fish catches) and spot-leaving decisions (Fig. 1C–E). The original study [7] analyzed 10 competitions; here we focus on a subset of five of these in which we elicited participants’ prior spatial beliefs before the competition started (see Quantifying Prior Beliefs and Fig. 1B). These five competitions took place on five subsequent days in March 2023 on five lakes in Northern Karelia, Finland. On each day, 51–56 highly experienced (median [SD] years of experience = 50 [13.7]; see also Table S2) competitive ice-fishers participated. Most participants took part repeatedly (number of unique participants = 62; mean number of days participated [SD] = 4.4 [1.19]; see also Table S3). All participants provided written informed consent prior to study participation.

### 2.2 Quantifying Prior Beliefs

To assess individuals’ prior beliefs about the resource distribution, each morning before the competition started we gave participants a topographical map of the lake-bed and asked them to mark the areas they would visit if they were foraging by themselves to maximize their catch (using a three-point confidence scale: red: would definitely visit; yellow: would very likely visit; green: would likely visit; with an upper limit of three areas per confidence level; Fig. 1B). Foragers were asked about locations they would visit if fishing alone to ensure that choices would not be affected by the anticipated competition at specific locations. The map’s contour lines (isoclines) showed depth gradients (see Figs. S1 and S2). Areas close to contour lines are areas where depth curves are bent and/or multiple depth curves are in close proximity, indicating the presence of underwater bays, islands or steep gradients. Anecdotal reports of ice-fishers suggest they use this feature, which we will denote roughness below, to identify promising areas, and previous work found that areas close to depth curves were indeed slightly more likely to be visited during comeptitions [7]. Maps were obtained from Retkikartta (https://www.retkikartta.fi/; last accessed on May 26th, 2025), a navigation software that many ice-fishers in Finland use to prepare for competitions.

Afterwards, the map surveys were scanned and each pixel was assigned a coordinate based on known coordinates of the map corners. To digitize the nominated areas, we subjected colored pixels to a DBSCAN [28] clustering algorithm to identify pixels belonging to the same nominated area. We then computed alphahulls (a more robust alternative to convex hulls) using the R package alphahull [29] around each identified cluster to obtain polygon shapes. To extract the confidence level for each area, we extracted the color of each area using DBSCAN applied to RGB values of each pixel. To ensure reliability of the RGB clustering method, colors were intensified using the software GIMP. A visual comparision of all digitally extracted areas and the original map surveys confirmed that this method accurately captured the shape and color of participants’ markings. See Fig. S3 for an illustration of the digitization procedure.

GPS coordinates for each of the isoclines displayed in the map surveys were obtained from the Finnish Land Survey (https://asiointi.maanmittauslaitos.fi/karttapaikka/tiedostopalvelu/maastotietokanta?lang=en), the Finnish Environment Institute (https://www.syke.fi/fi-FI/Avoin_tieto/Paikkatietoaineistot), and the Finnish Transport and Communications Agency (https://julkinen.traficom.fi/oskari/?lang=en; all last accessed February 4th, 2025).

### 2.3 Headcam and GPS Data Processing

The first author (A.S.) manually annotated all headcam data recorded during the competitions, extracting spot-leaving times and fish catches using the software Boris [30] (for details, see [7]; see also Fig. 1C–E). Spot-leaving times were defined as the duration between the time point at which the lure first entered the water and the time point at which the lure was taken out of the water for the last time. Fish catches were annotated at the time anglers lifted the fish out of the water. All annotated angling sequences were then assigned a GPS location by averaging all GPS fixes (1Hz frequency) during the angling event. To ensure data quality, missing data—defined based on the distribution of GPS error and the duration of obstructed video footage—were excluded from the analyses [7]. In particular, locations were excluded if the video recording was either obstructed for more than 30s during the start or end of an angling sequence, or for more than 10% of the angling time, if either the GPS noise at angling spots exceeded 5m or GPS was missing for more than 10% of the angling time, or if fish catches could not be determined due to video obstructions. All subsequent spots after the first excluded location were removed from analyses as the spot selection model (see below) predicts spot choices from temporally updated predictors.

### 2.4 Assessing Ecological and Social Characteristics of Prior Beliefs

#### 2.4.1 Area-Nomination Model

We first tested whether participants used the lake topography when nominating areas. Specifically, we tested the role of five topographical features that we hypothesized could inform their prior beliefs: (i) the distance to the starting point of the competition (displacement), (ii) the closeness (i.e., negative distance) to the competition boundary (boundary), (iii) the depth value of the closest isocline (depth), (iv) the closeness to the closest 0 depth isocline (shore), and (v) the closeness to the closest isocline (roughness) by comparing the actually nominated areas against simulated areas that could have been chosen by the individual. Features were computed using negative distances, such that higher values indicate a higher feature value (e.g., higher negative distances to the closest isocline indicates rougher areas). If topographical features are higher (lower) among the nominated compared to the respective simulated alternatives, this indicates that the topographical feature has a positive (negative) effect on the likelihood to nominate an area. A positive feature weight for depth, for example, indicates that individuals consistently nominate deeper areas compared to the simulated alternatives. Anecdotal reports and earlier results suggest that participants use roughness to locate underwater bays, islands or steep gradients in the lake bed which are thought to provide refuge for fish. Displacement was included to test whether individuals selected areas that were easier to reach, and boundary was included to test for effects on area nomination due to the shape of the arena.

For each nominated area *a*, let *A* ∈ *{*1, …, *A*^∗^*}* denote the index of the nominated area among a set of *A*^∗^ ∈ ℕ areas that were potentially available to the individual. For each nominated area, we simulated N = 50 counterfactual nominations by first rotating the area around its origin by an angle drawn randomly from a uniform distribution, and then shifting the nominated area by random values in *x*and *y*-direction on the lake surface (thus, *A*^∗^ = *N* + 1) (see also Fig. 2A–F).

**Figure 2.**
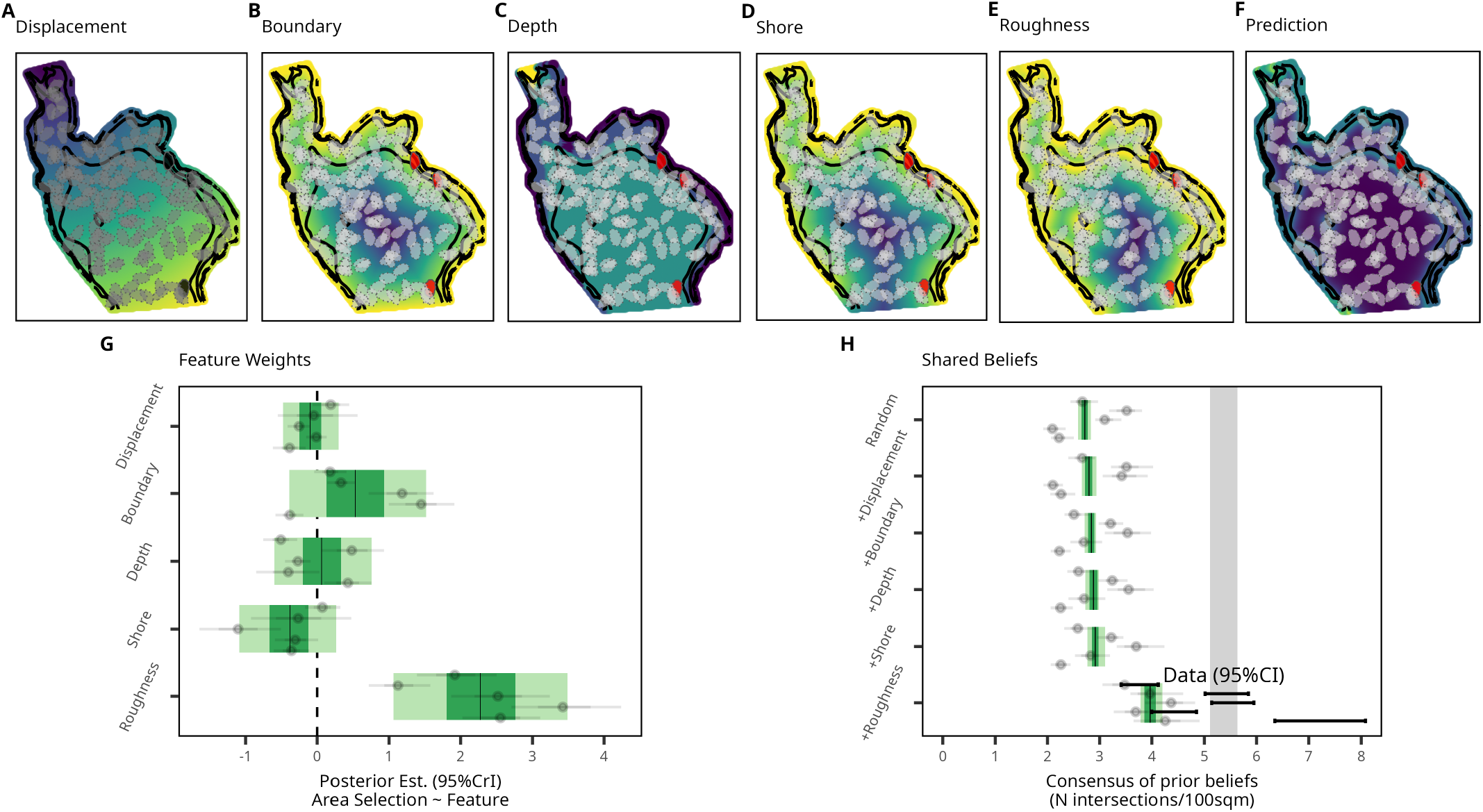
Assessing ecological and social characteristics of prior beliefs. (**A–E**) Visualization of the five lake-topographical features used to predict individuals’ prior beliefs. Nominated areas (red polygons) were compared with 50 counterfactual nominations (grey polygons) along five feature dimensions: (**A**) Displacement (the distance to the starting point, with lighter colors indicating higher displacement), (**B**) Boundary (the closeness to the arena boundary, with lighter colors indicating areas closer to the boundary), (**C**) Depth (the depth value of the closest depth curve, with lighter colors indicating deeper areas), (**D**) Shore (the closeness to the shoreline, with lighter colors indicating areas closer to the shore), and (**E**) Roughness (the closeness to the closest depth curve, with lighter colors indicating rougher areas). (**F**) Posterior predictions based on all five estimated model parameters. Lighter colors indicate higher probabilities of area nomination. (**G**) Feature weights predicting area nomination from the five spatial features (y-axis). Light (dark) green areas indicate 95% (66%) credible intervals for main effects pooled across the five lakes. Grey dots and intervals indicate posterior means and 95% credible intervals of lake-level effects. Numerical estimates can be found in Table S4. (**H**) Social consensus values predicted by models including no features (Random) to all five features. Average simulated consensus beliefs fall within the light (dark) green region in 95% (66%) of all simulation runs. Grey dots and intervals show 95% confidence intervals for each lake separately. Empirical consensus falls within the grey shaded area in 95% of all simulation runs and within the black error bars in 95% of simulation runs for each lake separately.

We model the nomination of each area as:

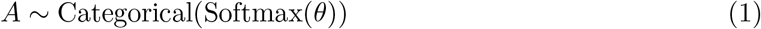

where 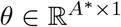 is a real-numbered vector of length *A*^∗^. Let *θ*_[*a*]_ denote the *a*-th entry of *θ*, then *θ*_[*a*]_ is given by

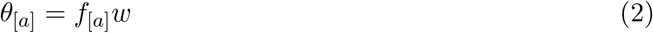

where *w* ∈ ℝ^*M ×*1^ is a real-numbered vector containing *M* parameters (i.e., feature weights) and *f*_[*a*]_ ∈ ℝ^1*×M*^ is a real-numbered row-vector containing *M* spatial features describing the candidate area with index *a*. The vector of spatial features comprises the ecological features described above. Values for each ecological feature were computed at a resolution of 100m^2^, and each area was assigned one value by averaging across all computed values for the respective area.

To account for the nested data structure (i.e., repeated observation of the same individuals across different lakes), we estimated feature weights *w* hierarchically with random offsets for participant, lake and foraging trip. Feature values were *z*-standardized across lakes before estimation to obtain standardized feature weights. Relative feature importance was additionally assessed using systematic model comparisons, fitting six different versions of the model: (i) a baseline model not containing any feature weights and selecting areas randomly, (ii) a model including only displacement as a predictor, (iii) a model additionally including boundary as a predictor, (iv) a model additionally including depth as a predictor, (v) a model additionally including shore as a predictor, and (vi) a model additionally including roughness as a predictor. Model performance was estimated by computing widely applicable information criteria (WAIC) based on expected pointwise log predictive densities (ELPD) relative to the best-fitting model using the R package loo [31]. Following [31], we concluded that models differed credibly in predictive performance if the difference in WAIC was larger than 4 times the standard error of the difference. To assess whether individuals used ecological features differently depending on confidence levels, we estimated an additional model including one feature weight for each confidence level.

#### 2.4.2 Consensus Estimation

To assess the degree to which prior beliefs were shared across individuals, we computed a measure of social consensus for each nominated area. We sampled 1 point per 100m^2^ from each nominated area and computed how many times that point fell into areas nominated by other individuals. For each area, the numbers across all sampled points were summed and the resulting number was divided by the number of sampled points, yielding a single consensus value approximating the spatial overlap across participants in intersections per 100m^2^. To assess whether consensus in the data was higher than expected by chance (i.e., whether the spatial overlap of nominations across participants was higher compared to the spatial overlap across random nominations), we repeated the same procedure with the N = 50 areas that were randomly shifted and rotated within the lake surface as part of estimating the effects of topographical features on area nominations outlined above. To assess whether social consensus was driven by individuals using the same ecological features to infer resource distribution, we also selected counterfactual areas as predicted by the posterior estimates of feature weights obtained from the participant data. That is, among the 50 random areas we selected areas according to the model predictions of the different versions of the area nomination model including different topographical features as predictors. Comparing those values to the empirical data allowed us to estimate the extent to which consensus was driven by these topographical features. To ensure reliability, we repeated this procedure N = 100 times for each actual area. We used the quantiles of the resulting distribution to compare values across conditions.

#### 2.4.3 Validity of Prior Beliefs

To estimate the validity of prior beliefs, that is, whether nominated areas were indeed more likely to be succesful, we conducted a Bayesian hierarchical zero-inflated Poisson regression predicting whether or not fish were caught at a angling spot (zero-inflation parameter) and the number of fish caught at each successful angling spot from whether or not the angling spot was nominated by the respective individual, and the number of times the angling spot fell within an area nominated by a competitor. To account for the hierarchical data structure, we added random slopes and intercepts for each lake, foraging trip and individual.

### 2.5 The Influence of Prior Beliefs on Foraging Behaviour

#### 2.5.1 Descriptive Analysis of Prior Belief Use

We obtained a descriptive measure of prior belief use by computing the fraction of angling spots within each trip that fell within the areas nominated by the respective individual. We defined a spot as falling within a nominated area if it was inside the boundaries or if the distance to the boundary was below a threshold, *ε*, of 100m. As a robustness check, *ε* was varied from 0 to 500m. Again, we constructed a control distribution by randomly shifting and rotating each foraging trip N = 50 times within the lake surface and computing the same outcome measure.

#### 2.5.2 Spot-Selection Model

We next assessed the relative importance of prior beliefs on spot selection compared with other, more dynamic, sources of information, namely, whether a person was successful or not in an area (personal information), and the number of other ice-fishers in an area (social information). Using a recently developed model framework [7], we predicted the choice of each subsequent angling spot from a set of temporally varying spatial features representing the information available to individuals at each point in time.

Let *S*_[*t*]_ ∈ *{*1, …, *S*^∗^*}* denote the index of a chosen angling spot at time *t* from a simulated set of available spots. For each choice, we assumed that *S*^∗^ ∈ ℕ spots were available to the individual. N = 50 available spots were simulated by randomly sampling a radius and angle from the full empirical distribution of step-lengths and turning angles between subsequent angling spot visits during all foraging trips (i.e., *S*^∗^ = *N* + 1). Then, the choice of each spot can be expressed as

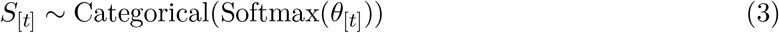

where 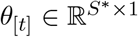 is a real-numbered vector of length *S*^∗^. The *s*-th entry of *θ*_[*t*]_, *θ*_[*t,s*]_ is computed as

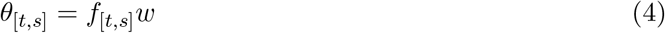

where *w* ∈ ℝ^*M ×*1^ is a real-numbered vector containing *M* parameters (i.e., feature weights) and *f*_[*t,s*]_ ∈ ℝ^1*×M*^ is a real-numbered row-vector containing *M* spatial features describing the spot with index *s*. The vector of spatial features contains available social, personal and prior information. Social information was computed as the kernel density of other competitors at the time the spot was chosen, personal information was computed as the kernel density of successful and unsuccessful angling spots visited by the individual, and prior information was computed as the closeness to the closest nominated area. Kernel densities were computed using a kernel function *K*.

Let *f*_[*t,s*,1]_ denote the first index of spatial features containing social information, then

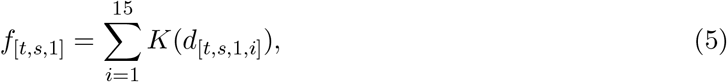

where *d*_[*t,s*,1,*i*]_ is the distance to the *i*−closest competitor. For computational tractability, *f*_[*t,s*,1]_ was only computed across the closest 15 competitors.

Let *f*_[*t,s*,2]_ denote the second index of spatial features containing the kernel density of successful spots, then

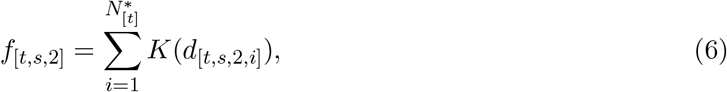

where *d*_[*t,s*,2,*i*]_ is the distance to the *i*−closest successful location. 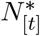 denotes the number of already encountered successful spots. For computational tractability, we set 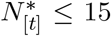. The computation was done analogically for the kernel density of unsuccessful angling spots.

The kernel function *K* determines the functional form of discounting of feature values with distance to competitors and angling spots. We chose a radial basis kernel such that

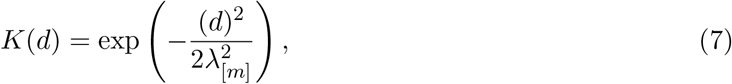

where *λ* ∈ ℝ^3*×*1^, a real valued vector of size 3. *λ*_[*m*]_ denotes the *m*-th element. *λ* determines the speed of spatial decay. By estimating *λ* as free parameters, we can investigate differences in spatial decay between features values. We assume, that spatial decay is the same for successful and unsuccessful spots.

To control for systematic differences in relocation distances that were not driven by information use, we added 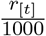 as a spatial feature, where *r*_[*t*]_ denotes the distance to the last known location in m. Additionally, we added 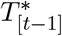 and 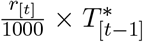 as spatial features, where 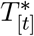 measures the time in the competition when starting to angle at the current spot. 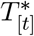 was normalized between 0 and 1, where 0 indicates the start of the competition and 1 indicates the end of the competition. Together, these features capture changes in relocation distances over time.

To estimate whether prior beliefs impacted the processing of social information (e.g., whether individuals were more/less attracted to nominated areas if other individuals were present) and personal information (e.g., whether individuals relocated less far from unsuccessful spots within nominated areas compared to outside nominated areas), we additionally estimated feature weights for the interaction between (i) the density of successful spots and the closeness to nominated areas, (ii) the density of unsuccessful spots and the closeness to nominated areas and (iii) social density and the closeness to nominated areas.

To estimate how information use varied across time, we additionally included feature weights for the interaction between (i) the density of successful spots and the time in the competition (normalized between 0 and 1), (ii) the density of unsuccessful spots and the time in the competition, (iii) social density and the time in the competition, and (iv) the closeness to the closest nominated area and time in the competition.

To assess the degree to which prior belief use differed as a function of the confidence level assigned to each area, we fitted an additional model including one feature weight for each level of confidence, respectively.

To assess feature importance, we conducted systematic model comparisons, fitting eight nested models: (i) a baseline model not containing any feature weights, assuming that individuals select spots randomly, (ii) a model including only locality and time in the competition as predictors, (iii) a model additionally including personal information, (iv) a model additionally including social information, (v) a model additionally including prior beliefs, (vi) a model additionally including the interaction effects (conditional weights) between prior beliefs and personal information, (vii) a model additionally including the interaction effects (conditional weights) between prior beliefs and social information, and (viii) a model additionally including the interaction effects (temporal weights) between spatial features and time in the competition. As before, model performance was estimated by computing differences in WAIC using the R package loo [31].

For all model parameters, we assumed weakly informative prior distributions, as detailed in [7]. Note that in [7] we showed that foragers selected areas closer to previously successful spots, further away from previously unsuccessful spots, closer to other competitors, and slightly closer to rougher areas. Here, we extend this model to study the role of prior beliefs on spot-selection decisions.

#### 2.5.3 Spot-Leaving Model

To study whether foragers remained longer (or shorter) in nominated areas compared to areas that were not nominated, we used the spot-leaving model described in [7]. This model predicts individuals’ spot-leaving decisions from a set of cues relating to the personal, social and prior information available to them at each point in time. That is, we predicted the probability that an individual leaves their foraging spot at time *t* from their immediate information on resource returns (i.e., time without a catch, *t*^∗^, and whether or not a fish has been caught, 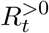). To assess the effect of the spatial features used to predict spot choices on spot-leaving decisions, we added the spatial features estimated in the spot-selection model. We denote these as *f*_Social_, *f*_Prior_, *f*_Successful_, *f*_Unsuccessful_. Again, we relied on systematic model comparisons to gauge the importance of each set of predictors.

Let each angling spot be characterized by a sequence of states *S*_[*t*]_ ∈ {0, 1} where *S*_[*t*]_ = 1 if the individual leaves at time *t* and 0 otherwise. Then,

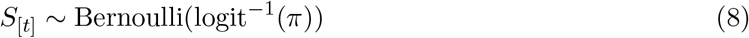

where *π* indicates the probability (on the logit scale) that the individual leaves the spot at time *t* and

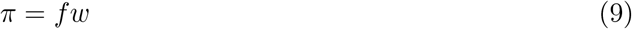

*f* is a row-vector containing features (i.e., 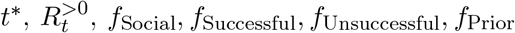) and *w* is a column-vector containing the respective feature weights.

Spatial features *f*_Social_, *f*_Successful_, *f*_Unsuccessful_ were computed using the mean posterior bandwidth estimates for each lake, *λ*_Social_, *λ*_Personal_, estimated in the spot-selection model detailed above. *f*_Successful_ and *f*_Unsuccessful_ were set to constant values for each angling spot, *f*_Social_ was computed for each timestep. All model parameters were estimated hierarchically with random offsets accounting for individual-, lake-, and trip-level variation in behaviour. For reasons of computational efficiency, we predicted leaving decisions at time intervals of Δ*t* = 10*s*. Note that in [7] we used this model to show that foragers stayed longer after catching the first fish at a location, were more likely to leave the longer they did not catch fish, and were more likely to stay in social areas. Here, we extend this model to study the role of prior beliefs on spot-leaving decisions.

### 2.6 Data Analysis

All reported statistical estimates are based on hierarchical Bayesian regression models controlling for the hierarchical data structure (i.e., foraging trips nested within participants and lakes). The area-nomination models, spot-selection models, and spot-leaving models were implemented in Stan [32] and run via the CMDstanR [33] interface using the statistical software R [34], with weakly informative priors for all model parameters [35]. For details on prior distributions, see [7]. We used within-chain parallelization across four chains with sufficient iterations to ensure an effective sample size of at least 200 for the main-level parameters. Chain convergence was assessed using the Gelman-Rubin statistic, 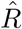, with values below or equal to 1.01 indicating chain convergence [36]. All other regression models were implemented using the R package brms [37] with default flat priors. We used the 95% posterior interval to discern credible from non-credible effects. Here we report the main effects from these models and hierarchical offsets for each lake. Standard deviations across all other hierarchical levels as well as numerical estimates for main effects are reported in the Supplemental Material.

## 3 Results

Across the five days, participants visited 10,091 angling spots. Of these, 1,610 spots (16.0%) were excluded (mainly due to headcams being blocked or out of position, see Table S1), leaving a total of 8,481 angling spots. This final dataset covers 251 foraging trips made by 62 unique individuals. On average, ice-fishers visited 33.8 spots per foraging trip, spent 205.1s at each spot, and 14.7% of spots were successful (i.e., one or more fish caught). On average, participants caught 1,255g of fish per foraging trip, with large variability between lakes (min = 483g, max = 2,012g). For a detailed sample description, see [7] and Table S2. In total, 808 areas were nominated in the map surveys, with a median area size of 0.02km^2^ (roughly the size of a 150m square). Participants nominated an average of 3.0 areas before each trip; 1.5 with high, 0.8 with medium and 0.7 with low confidence (Fig. S4).

### 3.1 Ecological and Social Chacteristics of Prior Beliefs

#### 3.1.1 Ecological Characteristics of Prior Beliefs

Comparing counterfactual and nominated areas, we found that lake-bed roughness was the strongest predictor of area nomination, with nominated areas being closer to depth curves than expected by chance (Fig. 2G). Displacement, the competition boundary, lake depth, or the lake shore did not predict area nomination (Fig. 2G). This pattern of results was corroborated by systematic model comparisons that showed that adding roughness led to substantial model improvement, while adding the other four features added little predictive accuracy (Fig. S5). Visual inspection of model simulations showed that the area-nomination model captured the distribution of area nominations well (Fig. S6). Feature weights did not differ across confidence levels (Fig. S7).

#### 3.1.2 Social Consensus in Prior Beliefs

We found that social consensus in area nomination was higher in the empirical data than expected by chance, suggesting that individuals shared similar prior beliefs about good fishing areas (Fig. 2H, bottom row). Simulating counterfactual nominations informed by the posterior estimates of the area-nomination model, we found that a large part of this consensus was explained by the common usage of the roughness feature to nominate areas (Fig. 2H, bottom row). The other topographical features did not substantially change the consensus estimates beyond the random baseline (Fig. 2H, top rows).

#### 3.1.3 Validity of Prior Beliefs

We did not find any credible association between prior nomination and foraging success, on either the individual or the collective level. In other words, individuals were not more likely to catch fish in areas they (or others) nominated than in areas they (or others) did not nominate (Fig. S8). Likewise, for spots where fish were caught, individuals did not catch more fish in nominated areas than in non-nominated areas (Fig. S8).

### 3.2 Prior Beliefs and Foraging Decisions

#### 3.2.1 Spot Selection

Foragers were more likely than chance to visit their nominated areas (Fig. 3A,B,C). In 66% of foraging trips, participants visited at least one of their nominated areas (chance expectation: 41%; Fig. 3A), and on average 37% percent of angling spots fell within a nominated area (chance expectation: 16%, Fig. 3B). This suggests that foragers relied at least partly on their prior beliefs when choosing spots. However, looking at visits to nominated areas for each foraging trip separately, we found that these effects were the result of a few individuals choosing most of their angling spots inside nominated areas: for 16% of foragers, more than 90% of spots were within nominated areas (chance expectation: 4%) (Fig. 3C). Descriptively, these affects are most pronounced for areas nominated with high confidence (Figs. S9 and S10).

**Figure 3.**
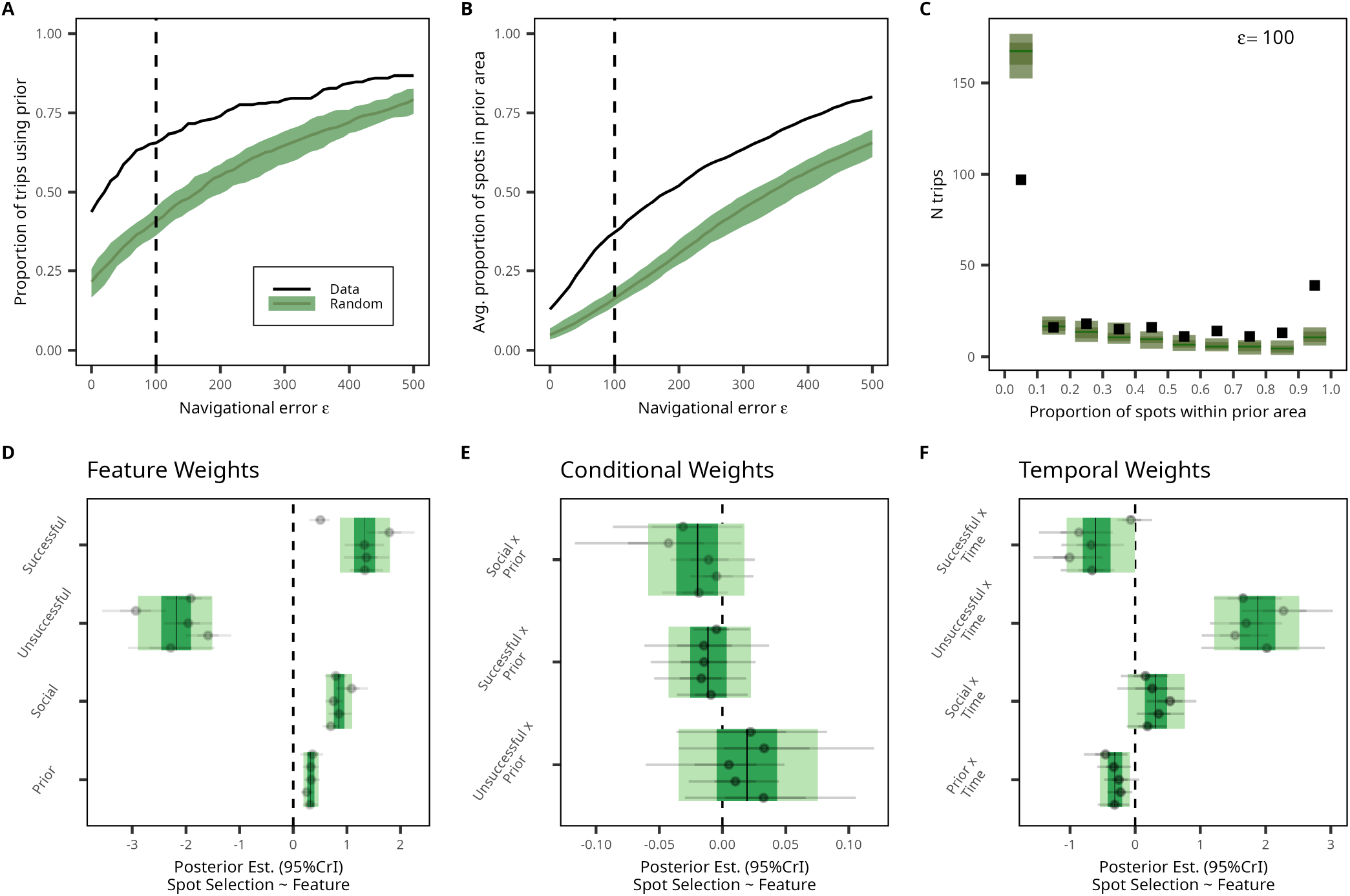
Prior beliefs and spot choices. (**A–C**) Descriptive analysis of prior belief use in spot choices. Proportion of foraging trips with at least one angling spot within a nominated area (**A**) and proportion of angling spots within a nominated area per foraging trip (**B**) accounting for potential error, *ε*, for empirical data (black line) and randomly distributed foraging trips; the green area indicates 95% of all simulation runs. The vertical dashed black line indicates navigational error of *ε* = 100m for which data is reported in the text. (**C**) Number of foraging trips as a function of the proportion of angling spots that fall within a nominated area (in steps of .1). Empirical data shown as black squares; simulated random foraging trips in green indicate 95% of all simulation runs. (**D–F**) Posterior feature weights, conditional weights (change in feature weights as a function of closeness to nominated areas), and temporal weights (change in feature weights across time in the competition; the start of the competition is coded as 0, the end of the competition as 1) of the spot-selection model. Displayed are standardized mean posterior estimates (thin black lines). The light (dark) green areas show the 95% (66%) credible intervals. Grey dots and intervals show posterior means and 95% credible intervals for each lake-level effect. Numerical estimates can be found in Table S5.

Turning to the spot-selection model, we found that (controlling for personal and social information use) prior beliefs had a credible, but small, effect on spot choices, with individuals selecting spots slightly closer to their nominated areas than expected by chance (Fig. 3D). In line with the descriptive measure of prior belief use, we find that the effect of prior beliefs on spot choices was strongest for the high confidence category (Fig. S11A). The difference in model estimates between different confidence levels was, however, not credibly different from 0 (Fig. S11B). As reported before in the larger sample, personal and social information had strong effects on spot choices [7], and these effects clearly outweighed prior beliefs. The effects of personal or social information were not altered by prior beliefs (Fig. 3E), as indicated by interaction terms that were not credibly different from 0. Looking at temporal variation in feature use we found that individuals were less likely to avoid unsuccessful areas, and were less strongly attracted to successful spots closer to the end of the competitions (Fig. 3F). Importantly, results also show that individuals were less likely to use prior beliefs towards the end of the competition compared to the beginning of the competition, suggesting that individuals update their prior beliefs throughout the competition (Fig. 3F). Generally, however, systematic model comparisons showed that including prior beliefs in the model did not credibly improve predictive performance beyond personal and social information (Fig. S12), indicating that, relative to the other features, prior beliefs played only a minor role for the decision where to search for resources. Providing support for the validity of the spot-selection model to capture individuals’ tendencies to rely on their prior beliefs, we found that the model estimates of prior belief use were strongly correlated with the descriptive measure of prior beliefs (Fig. S13).

In line with the fact that spots within nominated areas were not more successful than those outside of nominated areas, we found that use of prior beliefs for spot selection was not associated with either foraging success or percentage of successful spot visits (see Tables S6 and S7 for full model results).

#### 3.2.2 Prior Belief Use and Spatial Exploration

To better understand how search strategies differed between individuals who did and did not rely on prior beliefs, we correlated the trip-level posterior means of prior belief use with trip-level posterior means of attraction to successful spots and the avoidance of unsuccessful spots. We found that individuals who relied more strongly on prior beliefs relocated less far from unsuccessful spots (*b*[95%*CrI*] = −0.78[−0.89, −0.68]), but were equally likely to be attracted to succesful ones (*b*[95%*CrI*] = 0.05[−0.09, 0.18]) (Fig. 4A,B), suggesting a trade-off between sticking to one’s prior beliefs and avoiding losses. As a robustness check, we repeated this analysis, accounting for the full uncertainty in trip-level estimates of prior belief use and the avoidance of unsuccessful spots by extracting the posterior distribution of intercorrelations between trip-level estimates. Again, we found a negative correlation between the two estimates (*r*[95%*CrI*] = −.290[−.659, .184]). However, the 95% CrI included 0 due to the high uncertainty in trip-level estimates. Fig. 4C, D shows two foraging trips at opposite ends of this trade-off.

**Figure 4.**
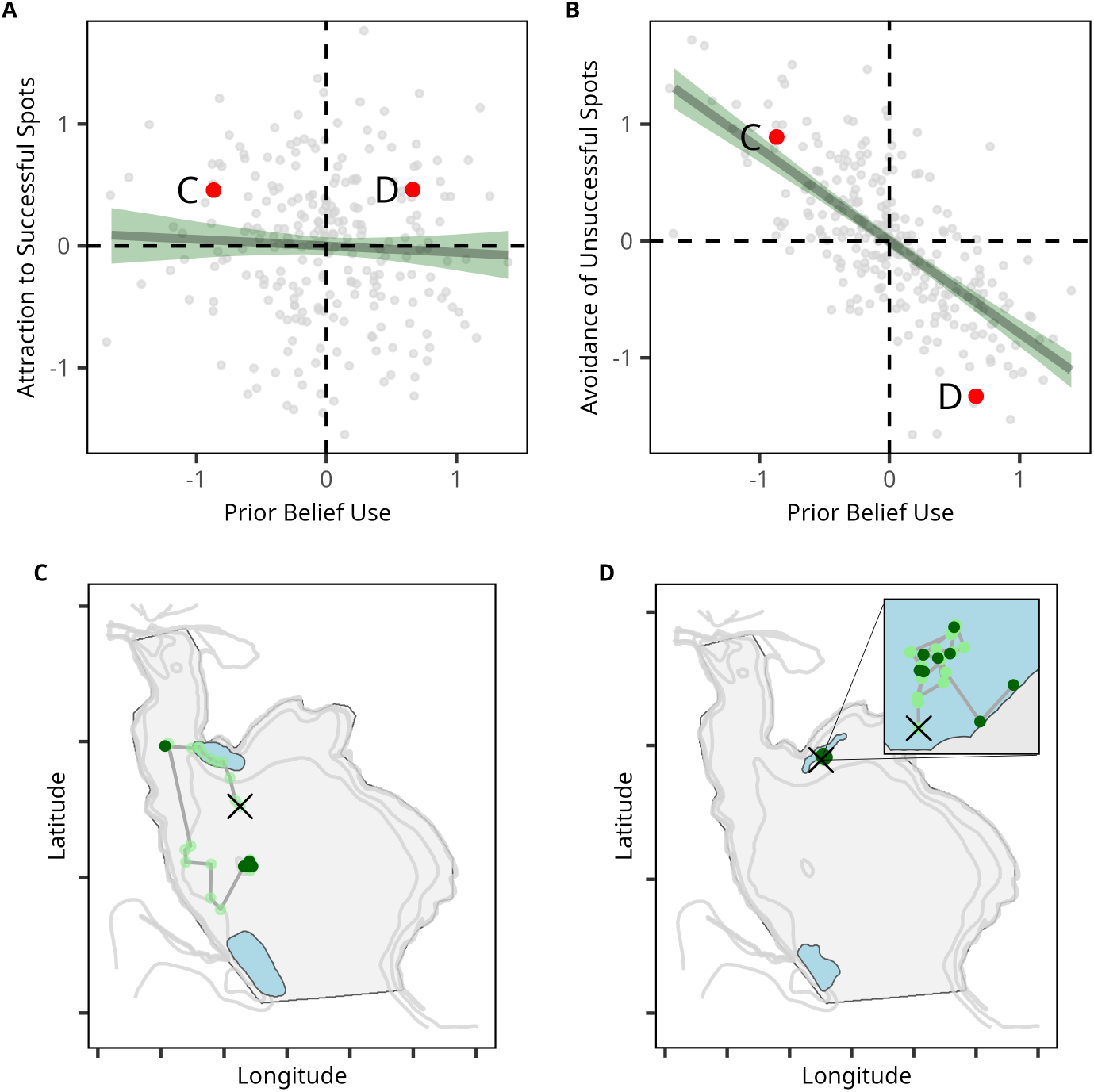
Prior beliefs and spatial exploration dynamics. (**A–B**) Association between trip-level estimates of prior belief use and attraction to successful spots (**A**) and avoidance of unsuccessful spots (**B**). Dark green lines (shaded regions) show the mean posterior prediction (95% credible intervals) of a linear regression analysis. Red dots highlight the two example foraging trips shown in (**C, D**). (**C**,**D**) Two foraging trips illustrating trips with varying degrees prior beliefs use and avoidance of unsuccessful spots. Crosses show the first angling spot, light green dots show unsuccessful angling spots, and dark green dots show successful spots (i.e., at least one fish caught). Blue shaded regions show the prior beliefs of the respective participant, ignoring confidence levels. (**C**) illustrates a participant not using their prior beliefs but exploring by relocating further from unsuccessful spots; (**D**) illustrates a participant sticking to their prior beliefs for the whole duration of the competition, thereby exploring limited space. Refer to Fig. S14 for randomly selected foraging trips with varying levels of prior beliefs use and avoidance of unsuccessful areas.

Trips with high prior belief use and trips low avoidance of unsuccessful areas explored only relatively small areas (Fig. S15). The negative effect of prior use on exploration disappeared, however, when controlling for individuals’ avoidance of unsuccessful areas (see Tables S8, S9, and S10), suggesting that reliance on prior beliefs reduces exploration via decreasing loss avoidance.

#### 3.2.3 Spot-Leaving Decisions

In line with our findings on spot selection, we did not find strong effects of prior beliefs on spot-leaving decisions. Instead, and as reported before in [7], information about resource return obtained directly at the angling spot was more important in predicting spot-leaving decisions than were spatial features (Fig. 5A and Fig. S16), with foragers spending slightly more time at spots in high social density areas (Fig. 5B). We also did not find that the effect of prior beliefs on spot-leaving decisions varied as a function of confidence levels (Fig. S17).

**Figure 5.**
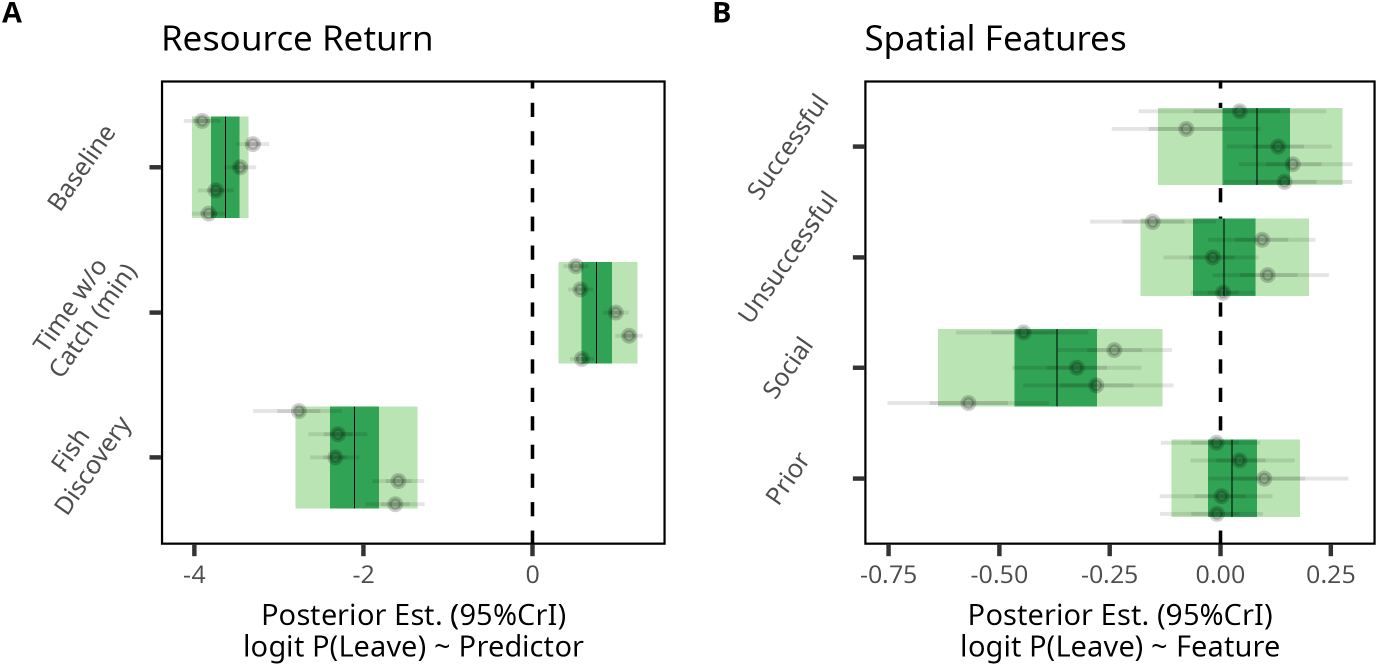
Spot-leaving decisions and prior beliefs. (**A**,**B**) Posterior estimates of the spot-leaving model. Displayed are standardized posterior estimates (thin black lines). The light (dark) green area shows the 95% (66%) credible intervals. Grey dots and intervals show posterior means and 95% credible intervals for each lake-level effect. Effects of (**A**) resource return rates (i.e., time without fish catches and fish discoveries on leaving decisions) and (**B**) the four spatial features on spot-leaving decisions. Numerical estimates can be found in Table S11.

Finally, using prior beliefs for spot-leaving decisions did not predict individuals’ overall foraging success (see Tables S12 and S13 for full model results).

## 4 Discussion

Foraging decisions are guided by complex information integration processes: Foragers need to evaluate dynamic information streams obtained through sampling the environment against their prior beliefs about the foraging ecology [2, 3, 38, 39]. Many social foraging models and laboratory studies highlight the importance of information sampling, learning and evidence accumulation during foraging [19, 40–42]. In real-world settings foragers possess rich cultural knowledge and experience about the foraging ecology, which might reduce the importance of information sampling and learning processes [27]. Here, we aimed to better understand the role of prior beliefs in social foraging, drawing on data from a large-scale field experiment in Finnish ice-fishing competitions. In addition to high-resolution GPS tracking and video annotation of fish catches and spot-leaving decisions, we obtained individuals’ prior beliefs about the resource distribution before each foraging trip, allowing us to quantify the relative importance of prior beliefs and dynamic information sampled during foraging on their foraging decisions.

Our analyses show that topographical features of the lake bed are used as a cue to inform prior beliefs about the presence of fish. These prior beliefs were also socially shared in our sample of participants, most likely due to individuals using the same topographical cues. This suggests that within this competitive fishing context, prior beliefs are to some extent informed by mental models that individuals hold about the relation between the topographical structure of the lakes and fish behaviour, rather than by accumulated prior experience made at specific locations. This has important implications for conceptualising information search in (social) foraging models: Rather than model-free accumulation of experience over time, foraging decisions might be better explained by mental models that guide information search [43]. Not accounting for such mental models might drastically affect the conclusions drawn—for example, about what constitutes adaptive decision-making [1, 44–46]. The large social consensus in what constitutes promising fishing areas also makes it likely that these mental models are reinforced through cultural transmission [47].

At the same time, we did not find evidence that individuals’ prior beliefs were systematically linked to foraging success. Both individually and collectively, prior beliefs were not predictive of successful spot visits or the number of caught fish at successful spots. Individuals relying more (less) on their prior beliefs for spot-selection and spot-leaving decisions were also not more (less) successful. These results are in line with our earlier findings showing that dynamically sampled information was predictive of successful spot visits, but that overall foraging success remains a highly stochastic outcome in fishing competitions [7]. It could be that the mental models individuals hold are effective only across large periods of time and repetition. An alternative explanation could be that the maps we provided were not at high enough resolution to draw reliable inferences about resource abundance. Indeed, the average size of nominated areas was very large, roughly comparable to a square of size 150m. A higher resolution of the topographical structure might result in more precise prior beliefs that might then also predict short-term foraging success.

While individuals to some extent used their prior beliefs to navigate the foraging landscape, they were quickly outweighed by more dynamic sources of information that provided cues about the resource distribution with higher spatial and temporal precision. Statistical decision theory predicts that in unpredictable or unstable environments the benefits of sampling novel information outweigh its costs, such that adaptive decision-makers should indeed rely less on prior beliefs [2] and use other sources of information such as social information to guide decision-making [48, 49]. In environments where conditions are generally stable across time, individuals might rely more strongly on prior beliefs and accumulated experience [50]. Experimental evidence, for instance, shows that in line with these assumptions humans tune their planning efforts to the environmental volatility with reduced planning in environments where outcomes are highly uncertain [51]. In the case of ice-fishing, we have found that capture likelihoods at a location decrease across time, suggesting that only the most recent personal and social information are reliable sources to locate good fishing spots [7, 13], in which case individuals’ foraging strategies would be well-tuned to the environment. Further research is needed to probe the role of prior beliefs systematically across foraging settings with varying degrees of ecological uncertainty.

Taken together, this study provides insights into the critical role of information sampling relative to prior beliefs during social foraging in stochastic and unpredictable environments. Spatial expectations about the resource distribution are, however, just one of the prior assumptions that individuals might hold. They may be less important drivers of decision-making than other ecological expectations and beliefs, such as the relation between predation and fish behaviour, the role of lighting conditions, the effects of local competition, the distribution of fish sizes, or the average size of fish schools. In our earlier work [7], for example, we found that giving up times are relatively stable within participants, but increase across lakes as resource abundance decreases, in line with predictions from optimal foraging theory [52, 53]. This suggests that prior beliefs about the average density of fish schools, and the likelihood to encounter them might affect spot-leaving decisions, while beliefs about the exact location of fish schools might not. Further research is needed to form a clearer understanding of the prior beliefs individuals hold about the resources and how these beliefs influence their foraging decisions.

## 5 Acknowledgements

### 5.1 Ethics

This study was approved by the Institutional Review Board of the Max Planck Institute for Human Development (A 2023-03). Before participation, each participant signed an informed consent form. We have complied with all relevant ethical regulations regarding data protection.

### 5.2 Data accessibility

All datasets and software code necessary to reproduce the results reported in this article can be accessed at https://github.com/aschakowski/Priors.

### 5.3 Declaration of AI use

We have not used AI-assisted technologies in creating this article.

### 5.4 Authors’ contributions

Following the CRediT standard [54]. **Conceptualization:** A.S., D.D., R.K., P.T.N., M.M.K., R.H.J.M.K.; **Data curation:** A.S.; **Formal analysis:** A.S.; **Funding acquisition:** R.K., R.H.J.M.K.; **Investigation:** A.S., D.D., R.K., P.T.N., M.M.K., M.P., R.H.J.M.K.; **Methodology:** A.S., D.D., R.K., P.T.N., R.H.J.M.K.; **Project administration:** A.S., R.K., R.H.J.M.K.; **Resources:** R.K., R.H.M.J.K.; **Software:** A.S.; **Supervision:** D.D., R.K., R.H.J.M.K.; **Validation:** A.S., D.D., R.H.J.M.K.; **Visualization:** A.S.; **Writing - original draft:** A.S., D.D., R.H.J.M.K.; **Writing - reviewing and editing:** A.S., D.D., R.K., P.T.N., M.M.K., M.P., R.H.J.M.K.

### 5.6 Funding

This work was funded by the Max Planck Institute for Human Development, and the Deutsche Forschungsgemeinschaft (German Research Foundation) under project number KU 3369, and under Germany’s Excellence Strategy–EXC 2002/1 “Science of Intelligence”, project number 390523135. The funders had no role in study design, data collection and analysis, decision to publish, or preparation of the manuscript.

## 5.7 Acknowledgments

The authors thank Tomi Keinänen, Mika Issakainen, the North Karelian Sport Fishing Association (Pohjois-Karjalan Urheilukalastajat ry), Petra Siemers-Hering, and Katja Münz for invaluable help with organizing the competitions and Susannah Goss for editing the manuscript.

## Supplementary Materials

### Supplementary Figures

**Supplementary Figure S1.**
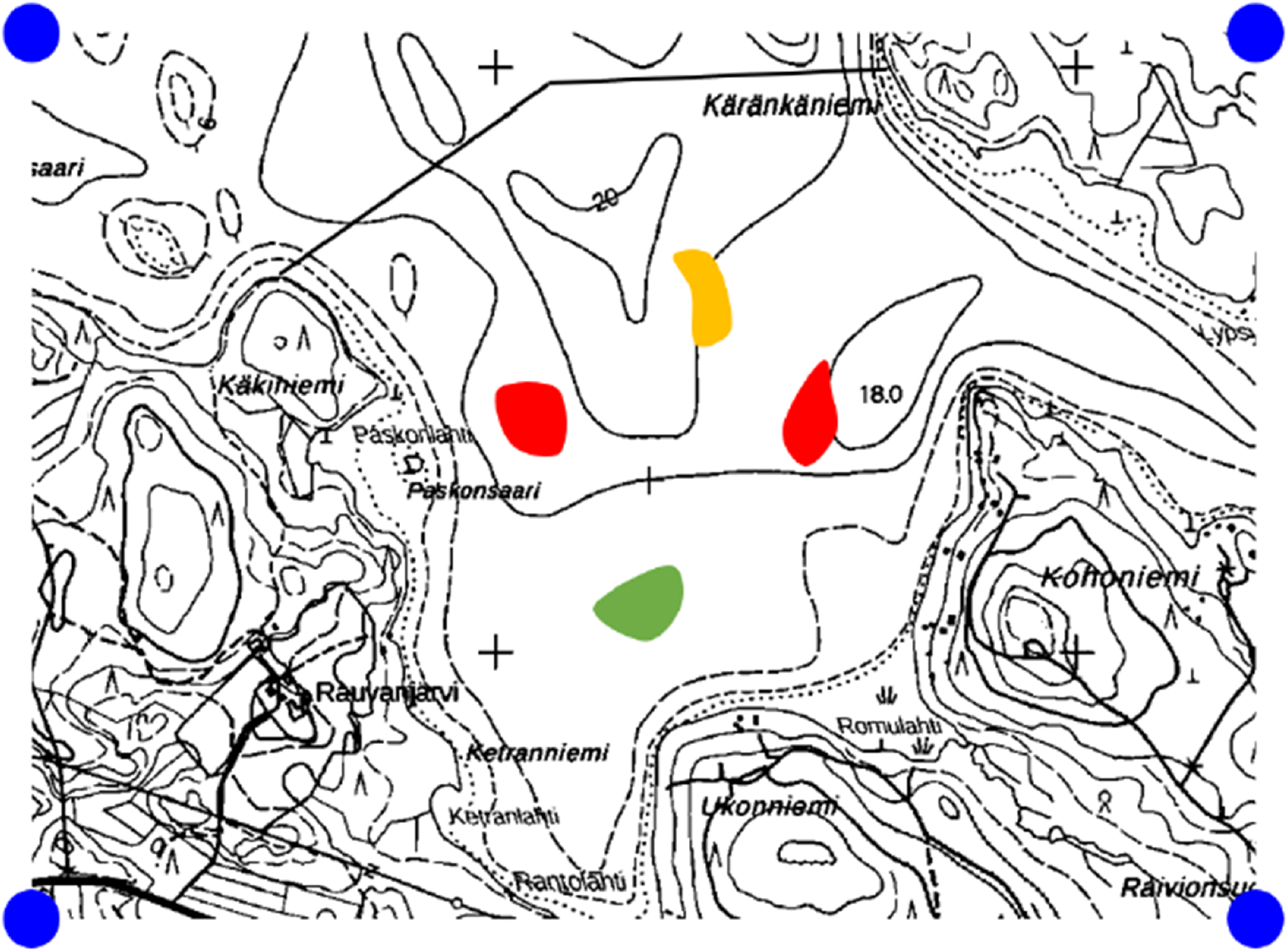
Assessment of prior beliefs: Instruction page. Participants were instructed to mark on a bathymetric map areas they would visit if foraging alone, using a three-point confidence scale. The survey was translated into Finnish by two native Finnish speakers (R.K. and M.P.).

**Supplementary Figure S2.**
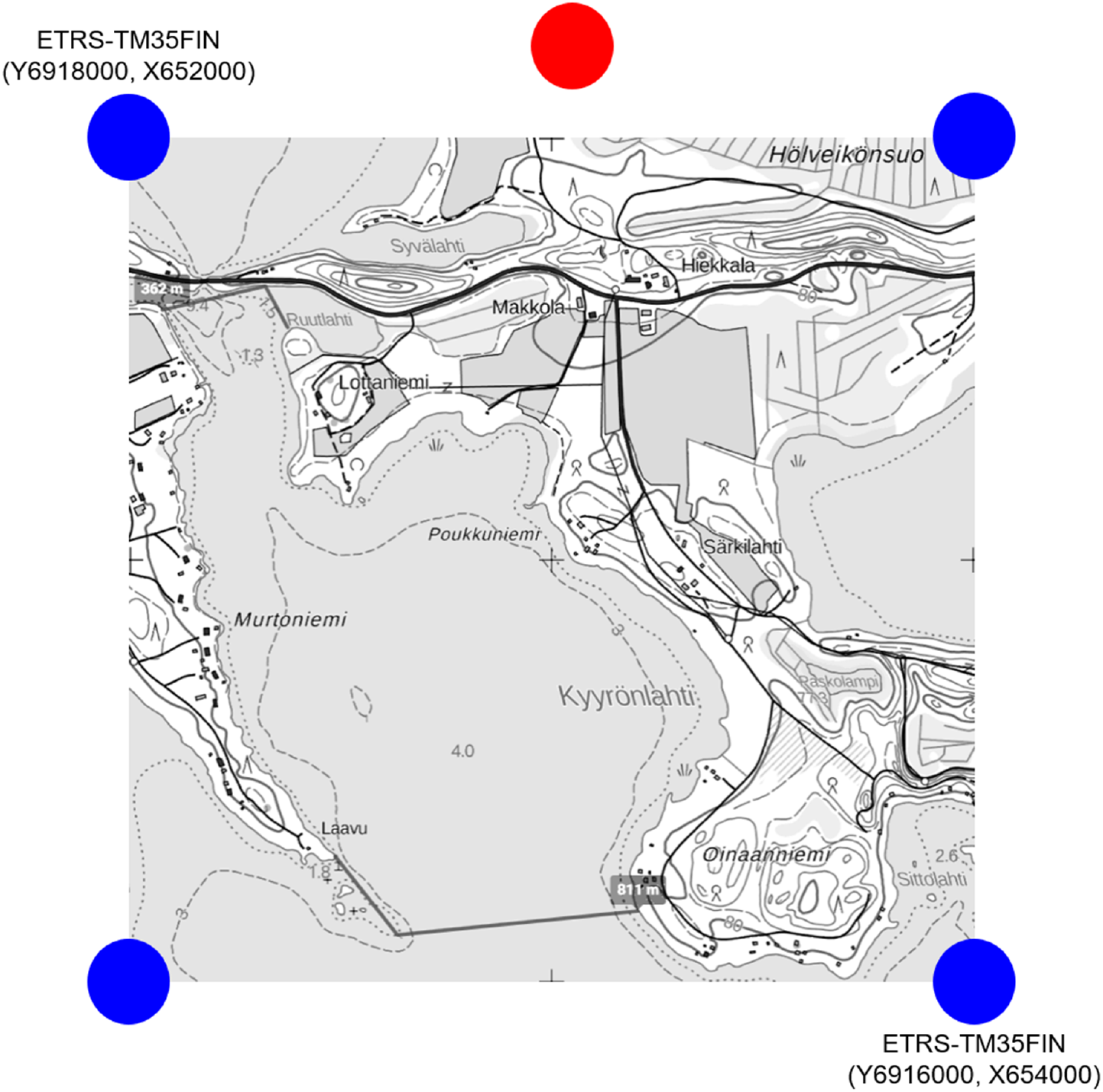
Assessment of prior beliefs: Response page. Participants marked the areas they would visit to maximize their catch on bathymetric maps before each competition. The blue dots mark the extent of the map excerpt, the red dot marks the top of the map. These indicators were later used to rotate scanned pages and to assign coordinates to the digitized marked areas. Note that the map used differed from the one shown in the instructions (see Figs. S1) to avoid any priming effect.

**Supplementary Figure S3.**
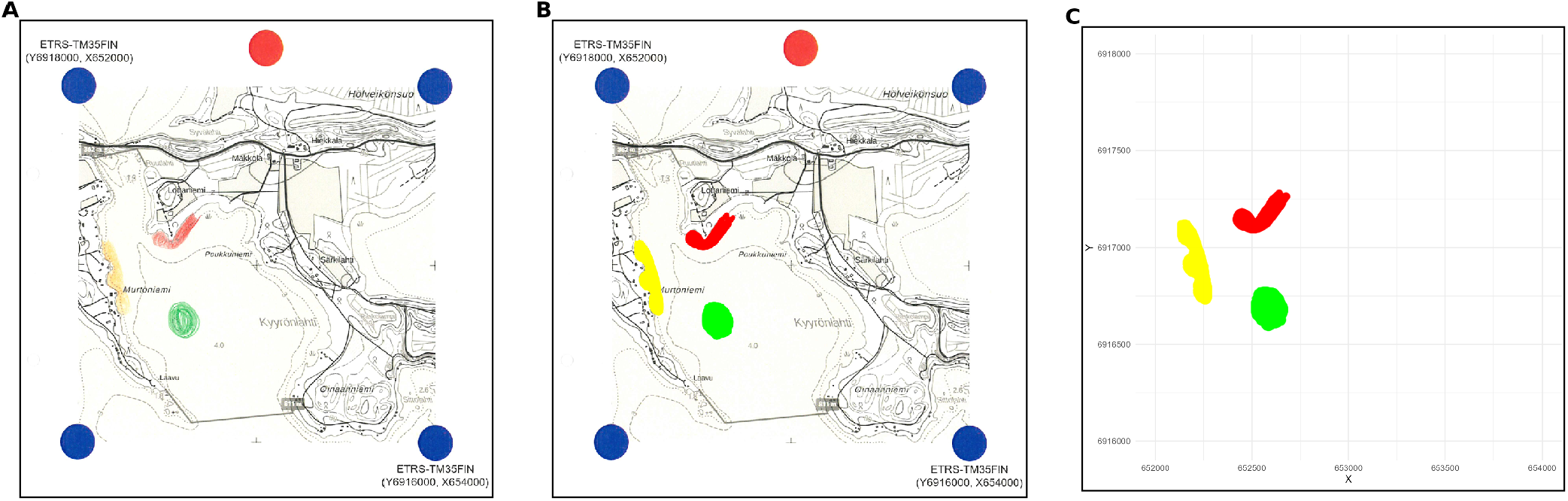
Digitization process for map surveys. (**A**) Example of a scanned map survey. (**B**) The same map survey with intensified colors to faciliate RGB clustering of confidence levels. (**C**) Digitized survey with map markings translated into alphahulls and assigned coordinates. Digitized areas were plotted next to the original surveys for each entry to ensure that the digitization procedure preserved shape, size and location of the marked areas.

**Supplementary Figure S4.**
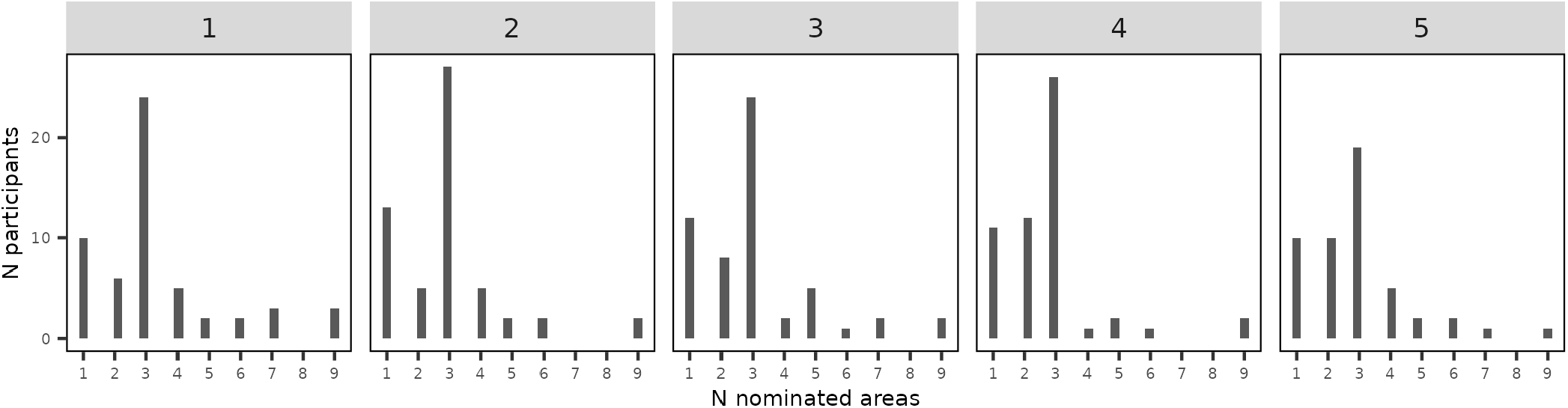
Distribution of area nominations by lake. Number of nominated areas (x-axis) per foraging trip per lake (1–5). Most participants nominated 1–3 areas per lake.

**Supplementary Figure S5.**
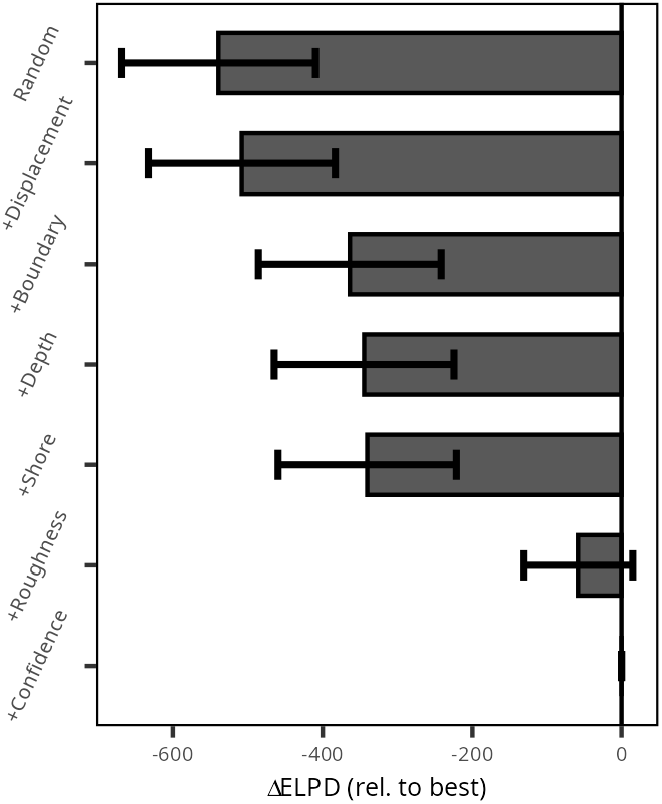
Model comparison area nomination model. Comparison of area nomination models assessing the predictive performance of five topographical features for predicting area nominations. Displayed are differences in widely applicable information criteria based on the pointwise expected log predictive densities (ΔELPD) relative to the best-fitting model (bottom row). Values closer to 0 indicate better model performance relative to the best-fitting model. The importance of each set of features can be assessed by comparing model performance to the most complex model not containing that set of features. Error bars indicate *±* 4*×* the standard error of the difference.

**Supplementary Figure S6.**
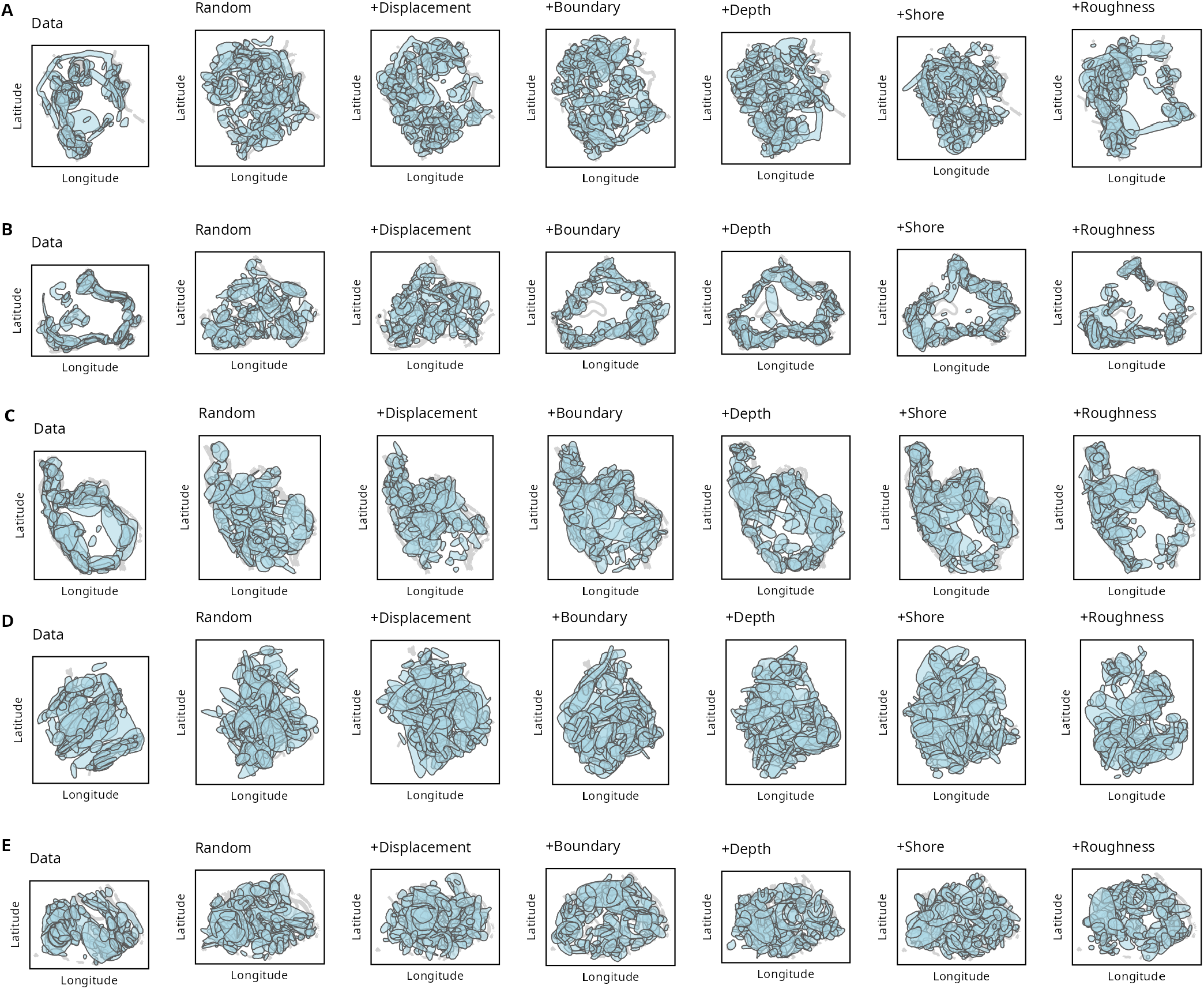
Simulation of area nominations predicted by the area-nomination models. (**A–E**) Displayed are all empirical nominated areas per lake (1st column; lakes in rows) and simulated area nominations from each version of the area-nomination model (2nd–7th column; lakes in rows) as light blue shaded regions on the five lake surfaces. Displayed is one randomly selected simulation run for each model. Visual inspections show that, in line with the results reported in Fig. 2F, adding the roughness feature increases overlap between area nominations across participants similar to the data.

**Supplementary Figure S7.**
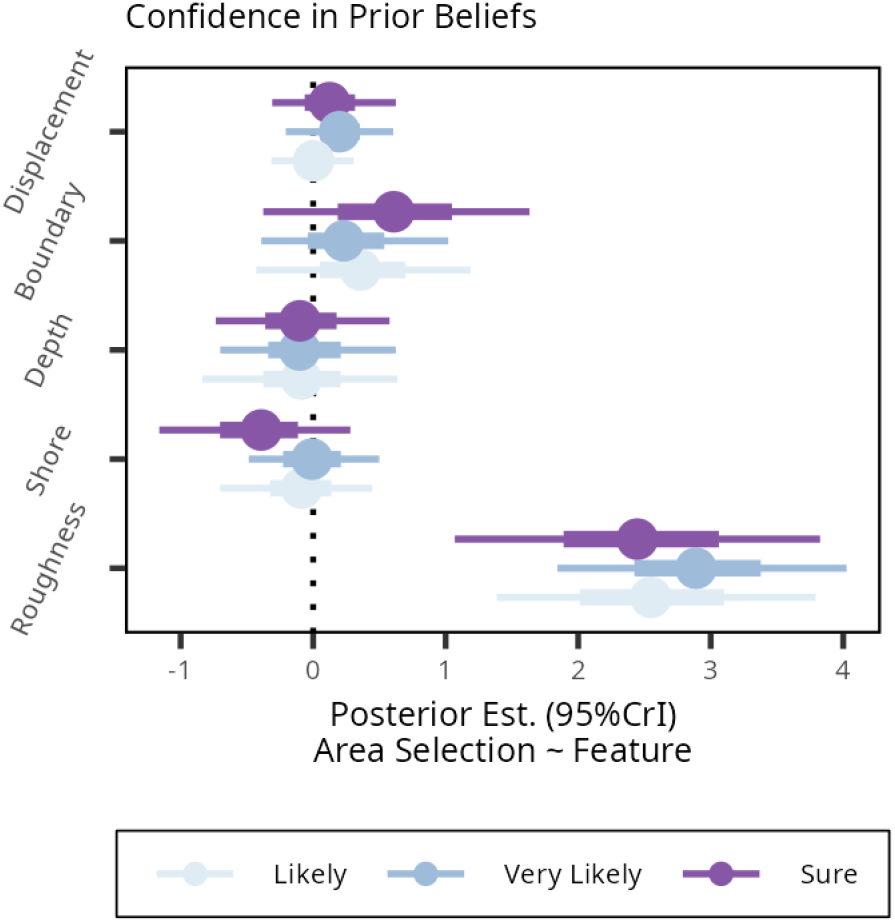
Effect of topographical features on area nomination as a function of confidence in prior beliefs. Dots display mean posterior estimates and (thin) lines show 66% (95%) credible intervals of the main effect of each topographical feature on area nomination as a function of its confidence level. Numerical estimates can be found in Table S14.

**Supplementary Figure S8.**
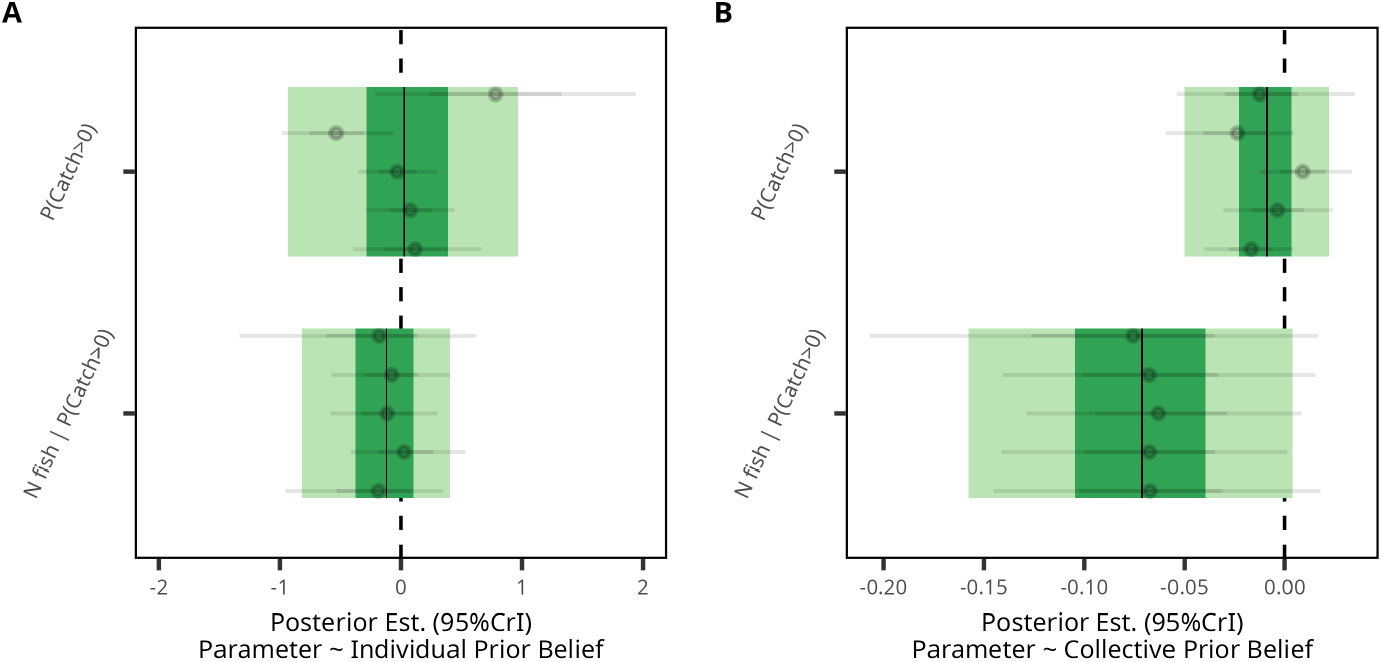
Validity of prior beliefs. (**A**,**B**) Posterior estimates predicting foraging success at each angling spot from prior beliefs. Foraging success was modeled using a zero-inflated Poisson-regression predicting from individual(i.e., whether or not a spot was contained in an area nominated by the same individual; **A**) and collective-level prior beliefs (i.e., the number of areas nominated by other individuals that the respective angling spot was part of; **B**) the probability of a catch (top row) and the number of fish caught (bottom row). Again, we assume that a location is within a nominated area, if the location is closer than *ε* = 100*m*. The dark (light) green areas show the 95% (66%) credible intervals. Grey dots and intervals show posterior means and 95% credible intervals for each lake-level effect. Numerical estimates are shown in Tables S15 and S16.

**Supplementary Figure S9.**
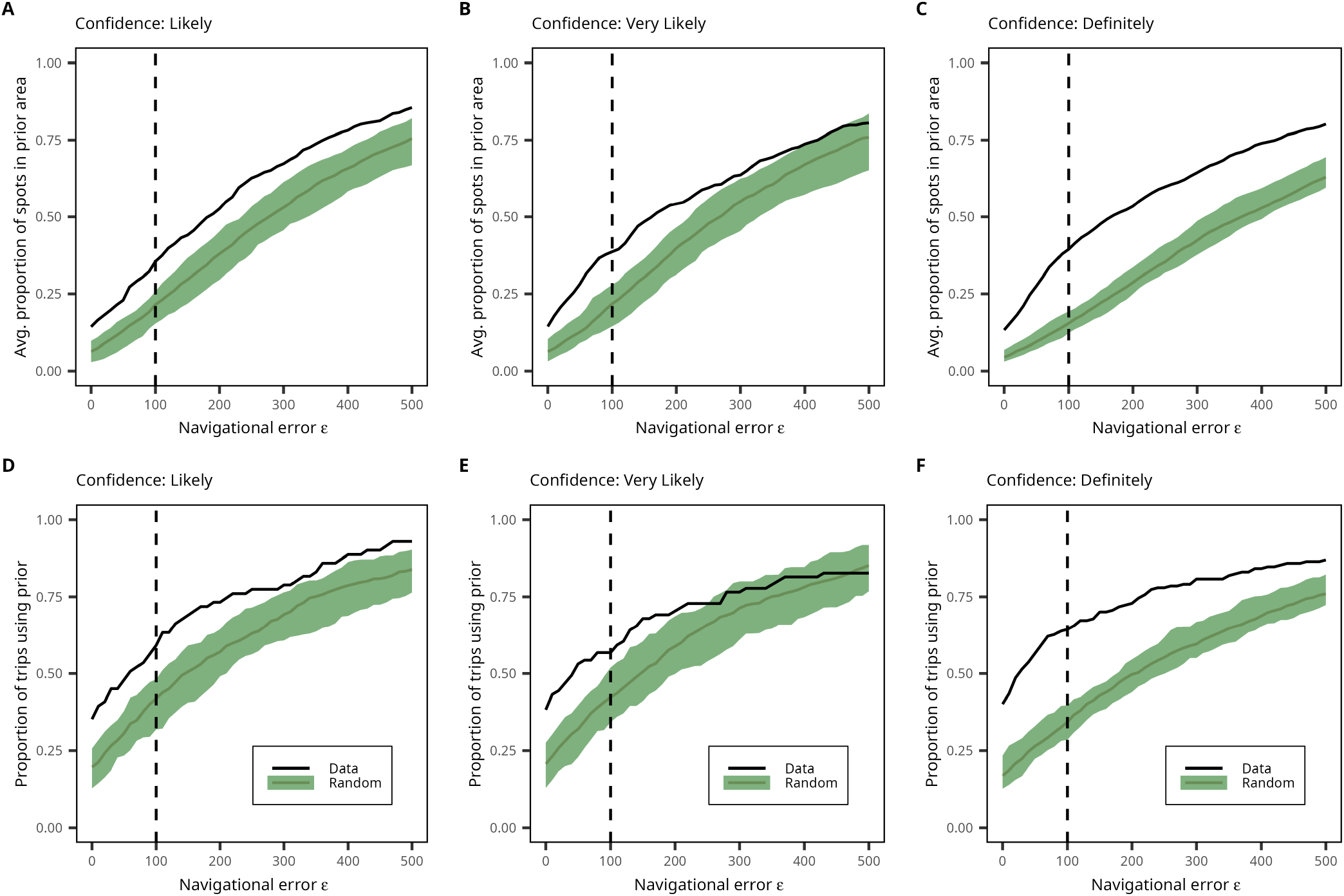
Descriptive measure of prior belief use by confidence levels. Descriptive analysis of prior belief use in spot choices. Proportion of angling spots within a nominated area per foraging trip across confidence levels. (**A–C**) and proportion of foraging trips with at least one angling spot within a nominated area (**D–F**) accounting for potential error, *ε*, for empirical data (black line) and randomly distributed foraging trips; the green area indicates 95% of all simulation runs. The vertical dashed black line indicates navigational error of *ε* = 100m.

**Supplementary Figure S10.**
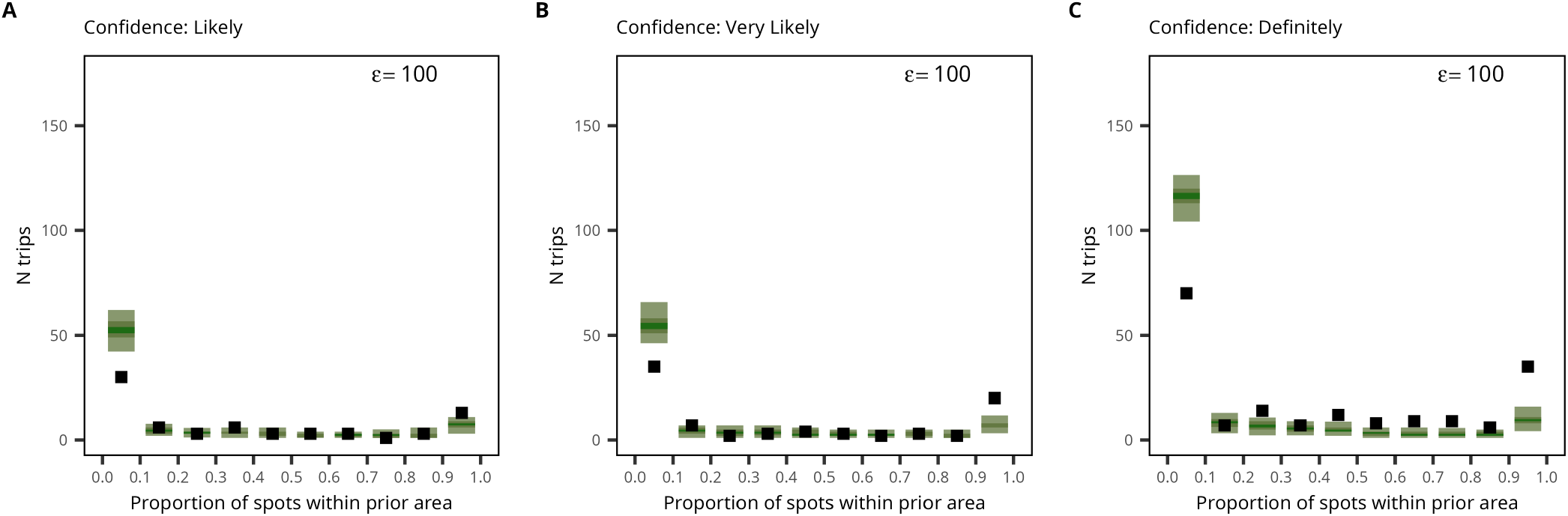
Descriptive measure of prior belief use by confidence levels. (**A–C**) Number of foraging trips as a function of the proportion of angling spots that fall within a nominated area (in steps of .1) split by confidence levels. Empirical data shown as black squares; simulated random foraging trips in green indicate 95% of all simulation runs.

**Supplementary Figure S11.**
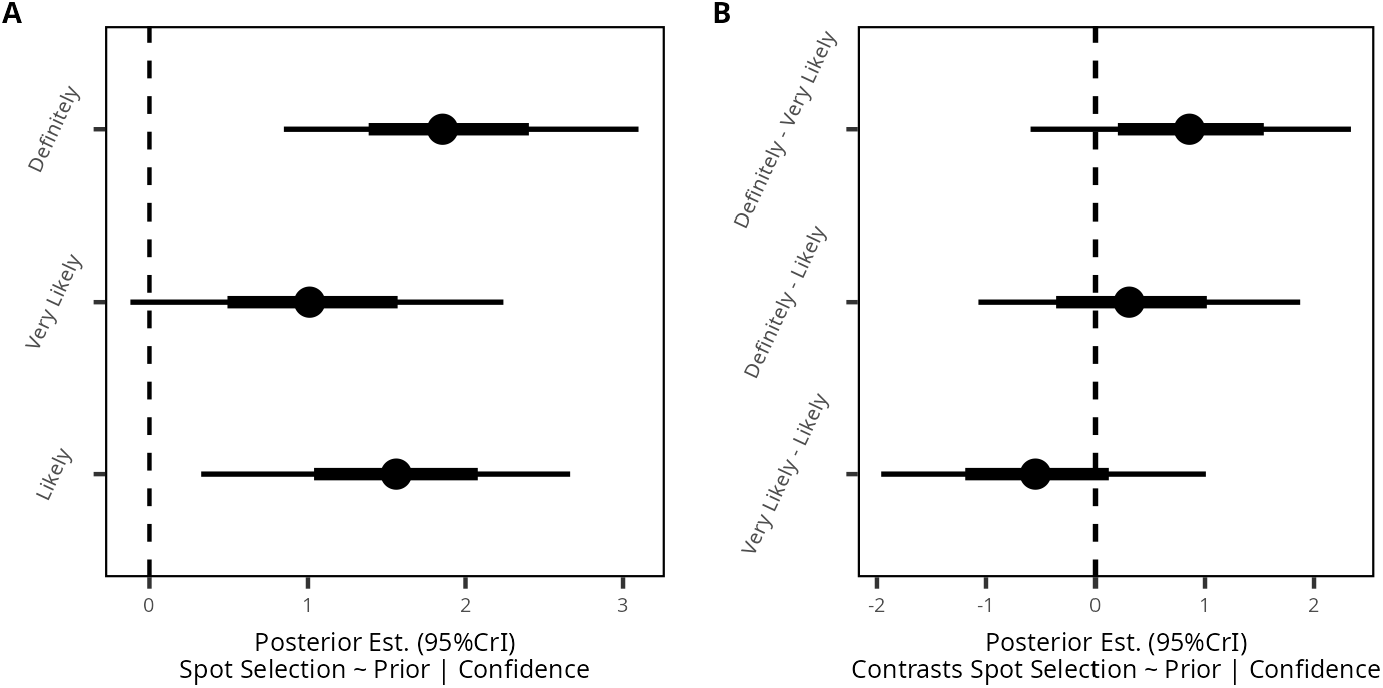
Effects of prior beliefs on spot selection by confidence levels. (**A**) Effects of prior beliefs on spot selection decisions split by confidence levels, and (**B**) posterior contrasts assessing the difference in estimates between the different confidence categories. Dots display mean posterior estimates and (thin) lines show 66% (95%) credible intervals. Numerical estimates are displayed in Table S17.

**Supplementary Figure S12.**
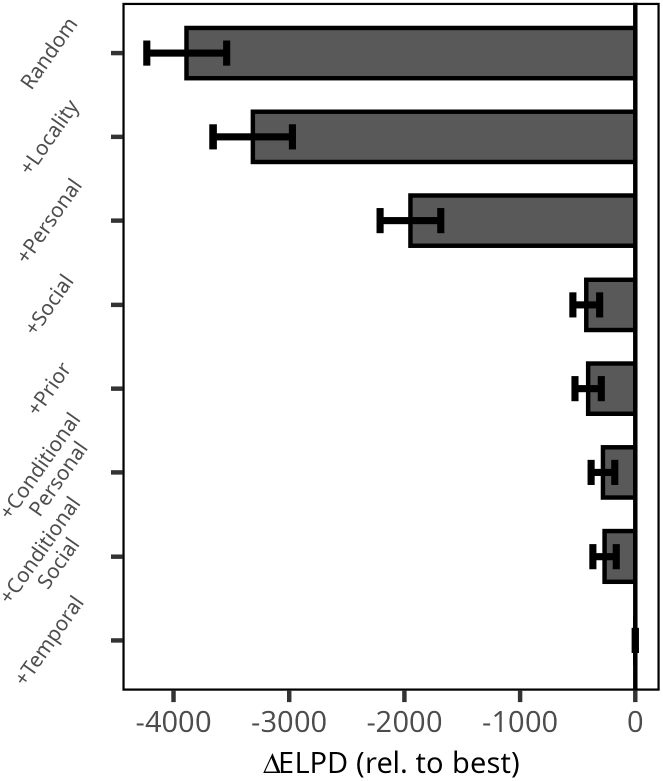
Model comparison spot selection model. Comparison of spot selection models assessing the predictive performance of the different spatial features for predicting spot choices. Displayed are differences in widely applicable information criteria based on the pointwise expected log predictive densities (ΔELPD) relative to the best-fitting model (bottom row). Values closer to 0 indicate better model performance relative to the best-fitting model. The importance of each set of features can be assessed by comparing model performance to the most complex model not containing that set of features. Error bars indicate *±* 4*×* the standard error of the difference.

**Supplementary Figure S13.**
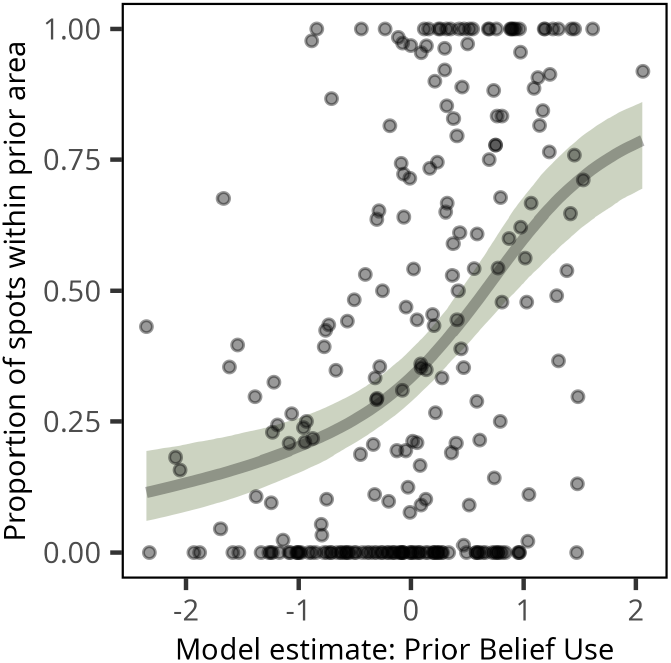
Convergence of descriptive and model-based measures of prior belief use. The relationship between a descriptive measure of prior belief use (i.e., the fraction of angling spots that were within a nominated area using an error margin of *ϵ* = 100m; see Methods) and the model-based estimates of prior belief use for each foraging trip. Black dots show data points and the green line (shaded region) shows posterior predictions (95% credible intervals) of a zero-inflated beta regression model predicting prior belief use from model estimates. Numerical estimates are displayed in Table S18.

**Supplementary Figure S14.**
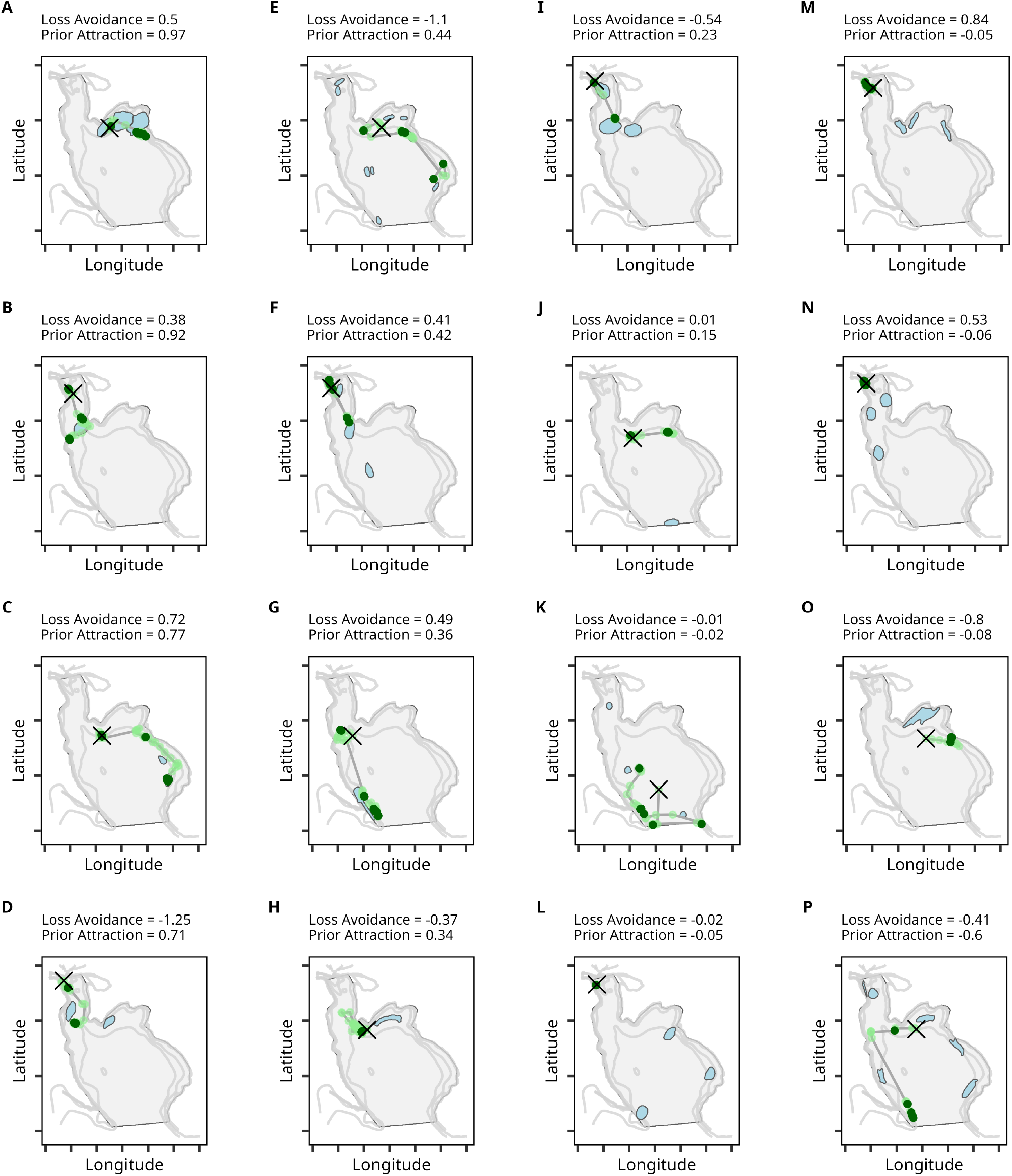
Example foraging trips illustrating prior belief use and avoidance of unsuccessful area. (**A–P**) Displayed are randomly selected foraging trips from one day illustrating the negative association between prior belief use (prior attraction) and avoidance of unsuccessful areas (loss avoidance). The black cross indicates the first angling spot of each trip, lightgreen dots indicate unsuccessful spots, darkgreen dots indicate successful spots, and lightblue areas indicate the prior beliefs of the respective individual. The examples are sorted according to the mean posterior estimate of prior attraction, with prior attraction decreasing from (**A**) to (**P**).

**Supplementary Figure S15.**
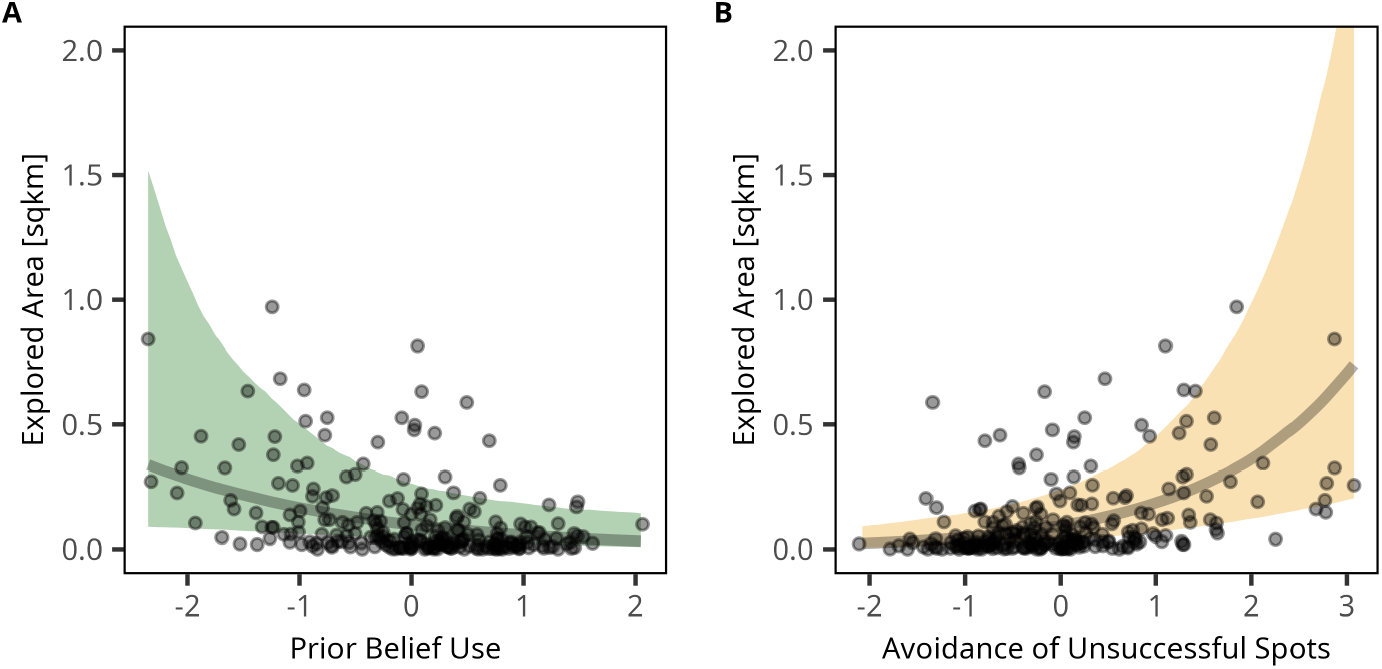
Prior belief use, loss avoidance and exploration. Displayed are posterior predictions of the average explored area as a function of mean posterior estimates of attraction to prior beliefs (**A**) and avoidance of losses (**B**). Black dots show foraging trips, lines show the mean prediction, shaded regions show 95% credible intervals. Numerical estimates are showns in Tables S8, S9, and S10.

**Supplementary Figure S16.**
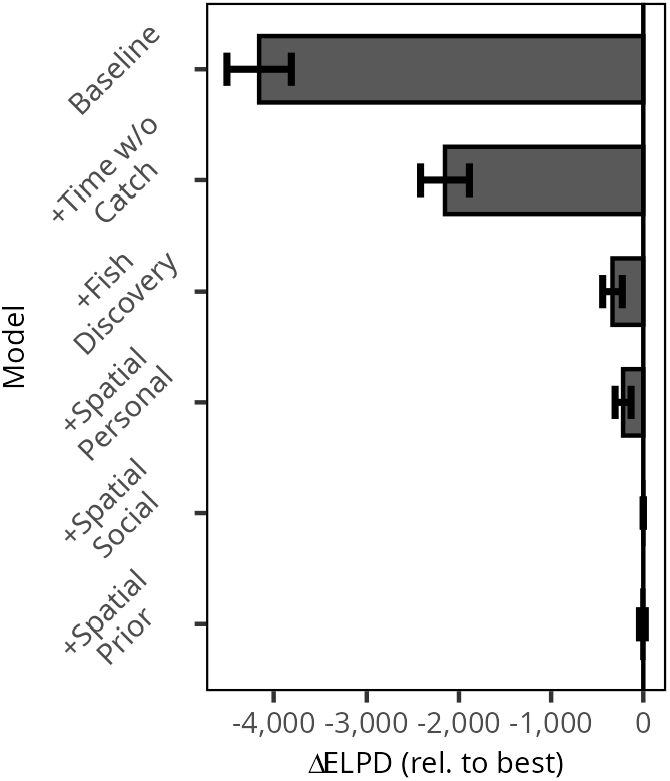
Model comparison spot leaving model. Comparison of spot leaving models assessing the predictive performance of the features for predicting spot leaving decisions. Displayed are differences in widely applicable information criteria based on the pointwise expected log predictive densities (ΔELPD) relative to the best-fitting model (second to bottom row). Values closer to 0 indicate better model performance relative to the best-fitting model. The importance of each set of features can be assessed by comparing model performance to the most complex model not containing that set of features. Error bars indicate *±* 4*×* the standard error of the difference.

**Supplementary Figure S17.**
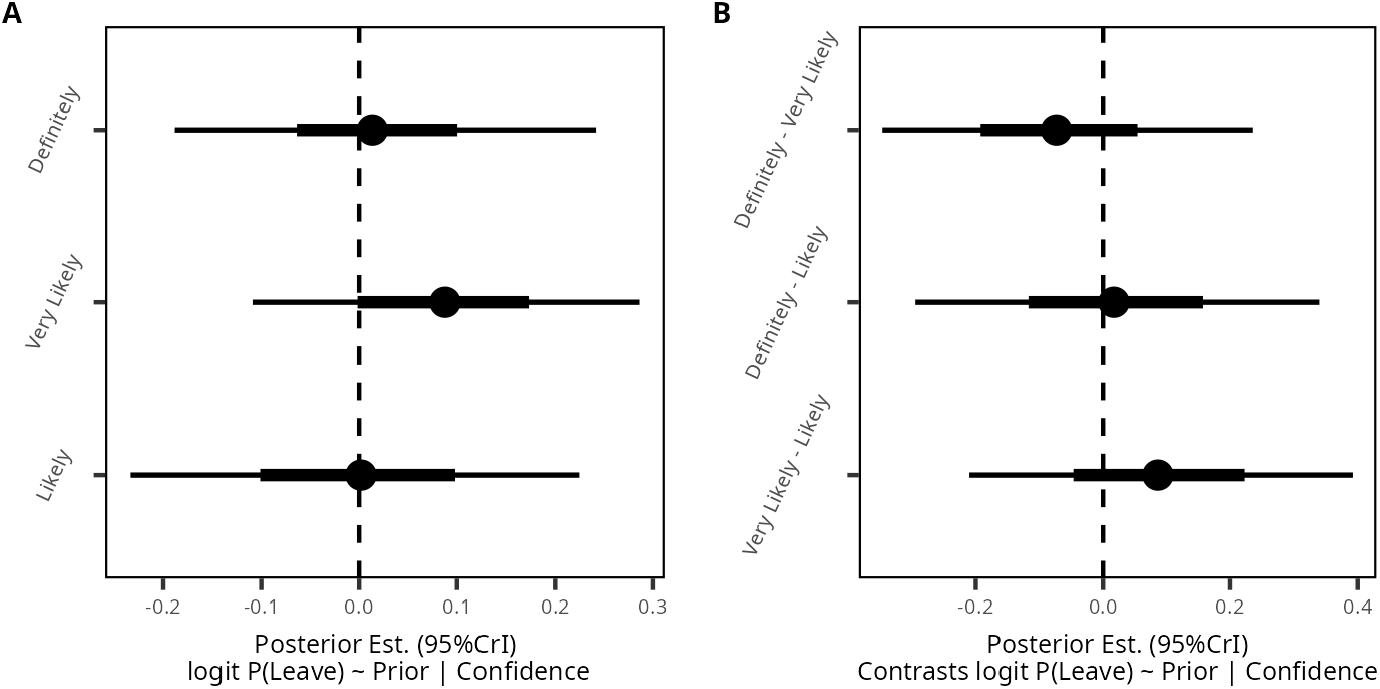
Effects of prior beliefs on spot-leaving decisions by confidence levels. (**A**) Effects of prior beliefs on spot leaving decisions split by confidence levels, and (**B**) posterior contrasts assessing the difference in estimates between the different confidence categories. Dots display mean posterior estimates and (thin) lines show 66% (95%) credible intervals. For numerical estimates, refer to Table S19.

### Supplementary Tables

**Supplementary Table S1.** Missing Data Angling Spots and Descriptive Statistics.

| day | 1 | 2 | 3 | 4 | 5 |
|---|---|---|---|---|---|
| Participants (N) | 55 | 56 | 56 | 55 | 51 |
| Missing watch/video recording (N) | 1 | 2 | 0 | 0 | 0 |
| Trips with only one spot (N) | 0 | 0 | 0 | 1 | 0 |
| Spots (N) | 2,363 | 1,737 | 2,033 | 2,055 | 1,903 |
| Start/end missing (N) | 26 | 11 | 15 | 13 | 0 |
| Video obstructed (N) | +6 | +15 | +7 | +8 | +0 |
| Noisy GPS (N) | +2 | +2 | +1 | +3 | +4 |
| Missing GPS (N) | +45 | +49 | +62 | +95 | +94 |
| Missing Catch (N) | +0 | +2 | +13 | +2 | +0 |
| Subsequent Spots (N) | +308 | +334 | +212 | +187 | +93 |
| Missing (N) | 387 | 413 | 310 | 309 | 191 |
| Missing (%) | 16.4 | 23.8 | 15.2 | 15.0 | 11.2 |
| Trips analyzed(N) | 52 | 46 | 53 | 51 | 49 |
| Mean spots per trip (N) | 43.8 | 32.2 | 36.3 | 38.1 | 37.3 |
| Median spots per trip (N) | 42.0 | 32.0 | 34.0 | 38.5 | 37.0 |
| Mean Missing per trip (N) | 7.17 | 7.65 | 5.54 | 5.72 | 3.75 |
| Median Missing per trip (N) | 0 | 0 | 0 | 0 | 0 |
| Trips with less than 5% missing (%) | 81.5 | 72.2 | 75.0 | 77.8 | 90.2 |
| Mean Successful spots per trip<sup>*</sup>(%) | 8.4 | 32.2 | 22.9 | 13.3 | 4.0 |
| Median Successful spots per trip (%) | 7.7 | 30.7 | 23.7 | 9.8 | 3.3 |
| Mean Angling Time successful spots<sup>*</sup> (s) | 687.0 | 705.5 | 852.9 | 1218.1 | 866.0 |
| Median Angling Time successful spots (s) | 500.0 | 594.8 | 745.7 | 625.0 | 683.3 |
| Mean Angling Time unsuccessful spots<sup>*</sup>(s) | 155.1 | 107.7 | 101.7 | 130.3 | 174.0 |
| Median Angling Time unsuccessful spots (s) | 151.2 | 94.8 | 97.8 | 123.1 | 170.5 |
| Mean fish catches per successful spot per trip<sup>*</sup>(N) | 2.6 | 14.0 | 15.5 | 12.4 | 2.2 |
| Median fish catches per successful spot per trip (N) | 1.5 | 11.5 | 10.4 | 5.8 | 1.3 |

Note. <sup>\*</sup>Results after exclusion of missing data.

**Supplementary Table S2.** Participant Characteristics per Competition.

| Lake | 1 | 2 | 3 | 4 | 5 |
|---|---|---|---|---|---|
| (N trips) | (55) | (56) | (56) | (55) | (51) |
| Mean catch (g) | 483 | 2,012 | 1,970 | 1,432 | 279 |
| SD catch (g) | (381) | (1,186) | (1,104) | (1,187) | (343) |
| Median catch (g) | 395 | 1834 | 1762 | 962 | 129 |
| Missing (N) | 0 | 0 | 0 | 0 | 0 |
| Mean age (y) | 61.8 | 61.2 | 61.2 | 61.3 | 61.1 |
| SD age (y) | 14.0 | 14.1 | 14.2 | 14.3 | 14.3 |
| Median age (y) | 65.0 | 65.0 | 65.0 | 65.0 | 64.0 |
| Missing (N) | 0 | 0 | 0 | 0 | 0 |
| Female (%) | 27.3 | 26.8 | 26.8 | 23.6 | 27.5 |
| Missing (N) | 0 | 0 | 0 | 0 | 0 |
| Mean experience (y) | 45.4 | 44.3 | 45.4 | 45.3 | 45.2 |
| SD experience (y) | 13.8 | 14.5 | 13.5 | 13.6 | 13.9 |
| Median experience (y) | 50.0 | 50.0 | 50.0 | 50.0 | 50.0 |
| Missing (N) | 4 | 3 | 4 | 3 | 3 |

Note. SD = standard deviation; y = years; g = grams.

**Supplementary Table S3.**
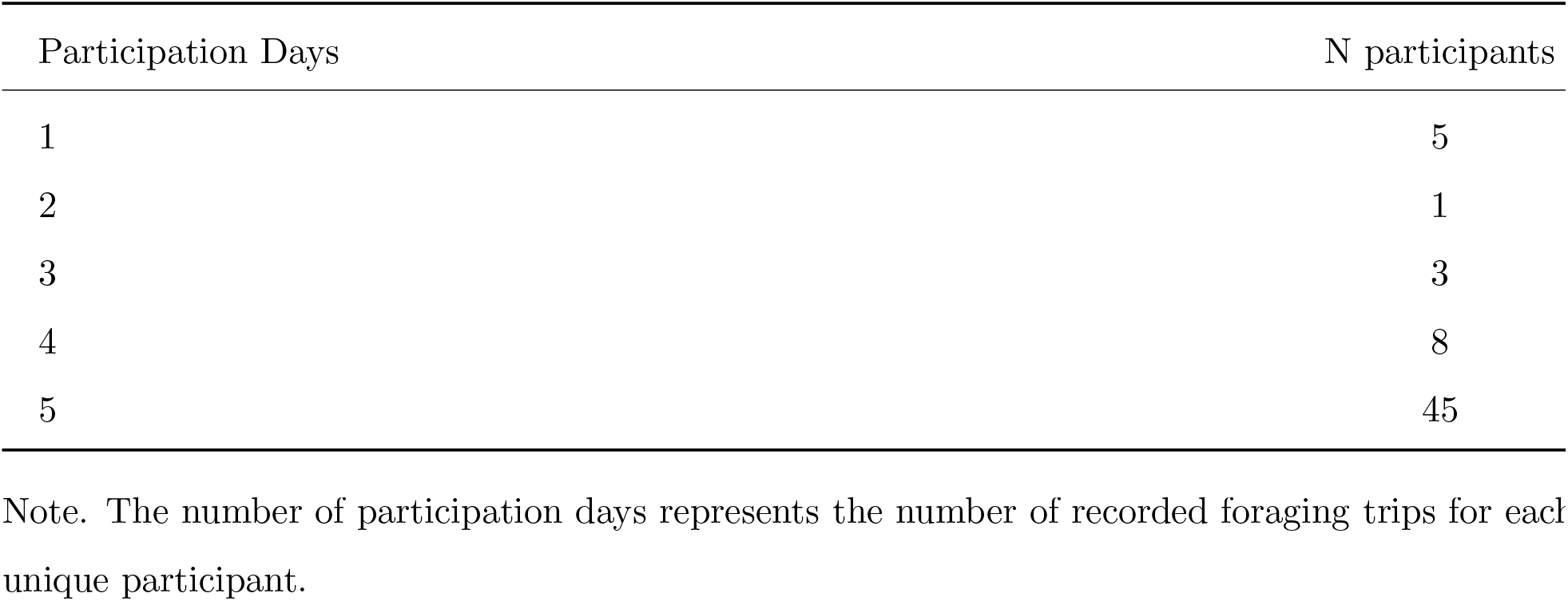
Participation Days per Participant.

| Participation Days | N participants |
|---|---|
| 1 | 5 |
| 2 | 1 |
| 3 | 3 |
| 4 | 8 |
| 5 | 45 |

Note. The number of participation days represents the number of recorded foraging trips for each unique participant.

**Supplementary Table S4.**
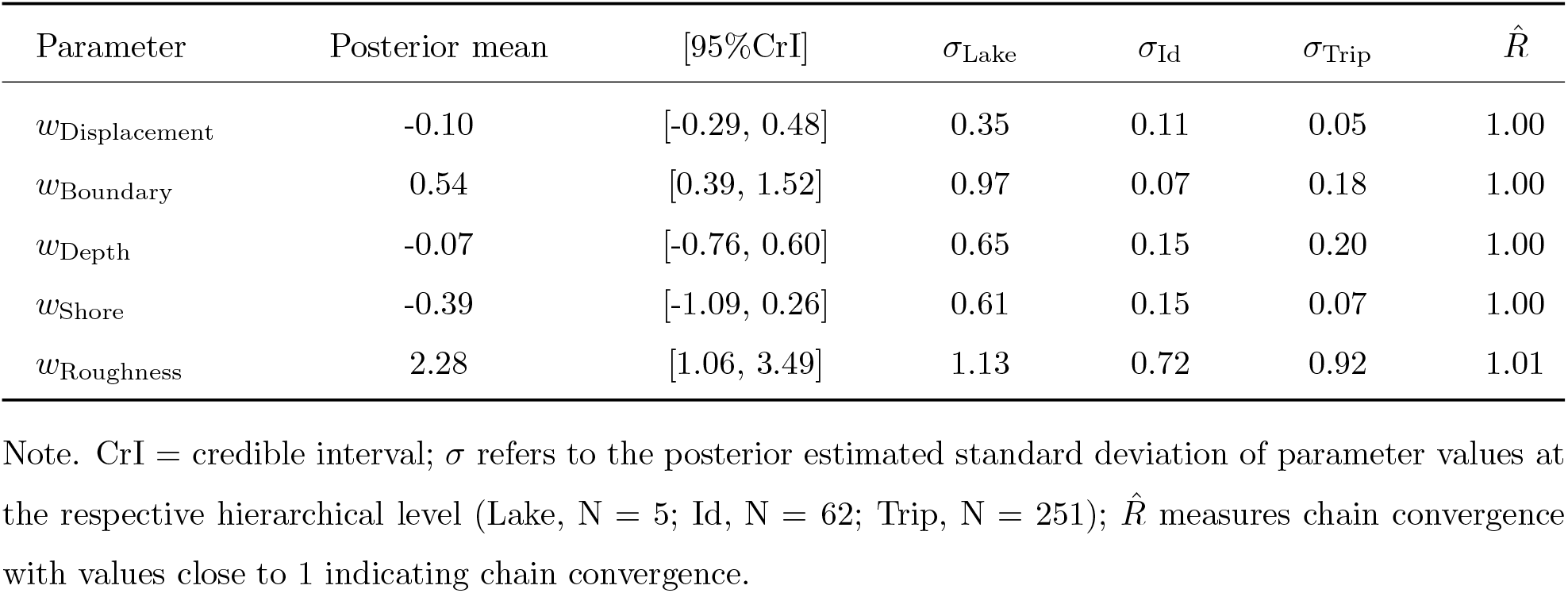
Standardized Posterior Estimates and Convergence Diagnostics of the AreaNomination Model.

| Parameter | Posterior mean | [95%CrI] | *σ*<sub>Lake</sub> | *σ*<sub>Id</sub> | *σ*<sub>Trip</sub> | $\hat{R}$ |
|---|---|---|---|---|---|---|
| *w*<sub>Displacement</sub> | -0.10 | [-0.29, 0.48] | 0.35 | 0.11 | 0.05 | 1.00 |
| *w*<sub>Boundary</sub> | 0.54 | [0.39, 1.52] | 0.97 | 0.07 | 0.18 | 1.00 |
| *w*<sub>Depth</sub> | -0.07 | [-0.76, 0.60] | 0.65 | 0.15 | 0.20 | 1.00 |
| *w*<sub>Shore</sub> | -0.39 | [-1.09, 0.26] | 0.61 | 0.15 | 0.07 | 1.00 |
| *w*<sub>Roughness</sub> | 2.28 | [1.06, 3.49] | 1.13 | 0.72 | 0.92 | 1.01 |

Note. CrI = credible interval; *σ* refers to the posterior estimated standard deviation of parameter values at the respective hierarchical level (Lake, N = 5; Id, N = 62; Trip, N = 251); $\hat{R}$ measures chain convergence with values close to 1 indicating chain convergence.

**Supplementary Table S5.**
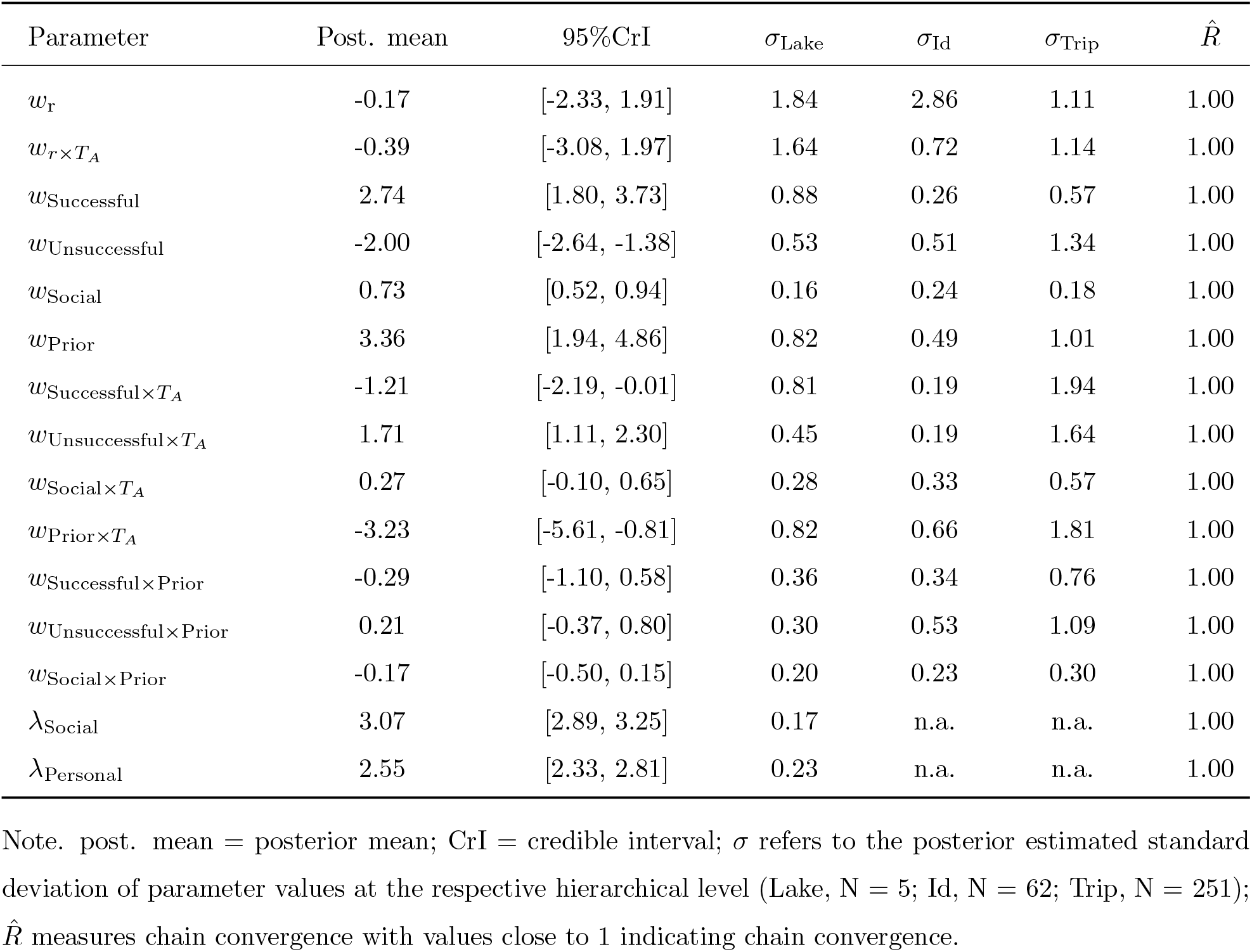
Unstandardized Posterior Estimates and Convergence Diagnostics Spot-Selection Decision Model.

| Parameter | Post. mean | 95%CrI | σ<sub>Lake</sub> | σ<sub>Id</sub> | σ<sub>Trip</sub> | $\hat{R}$ |
|---|---|---|---|---|---|---|
| *w*<sub>r</sub> | -0.17 | [-2.33, 1.91] | 1.84 | 2.86 | 1.11 | 1.00 |
| $w_{r\times T_A}$ | -0.39 | [-3.08, 1.97] | 1.64 | 0.72 | 1.14 | 1.00 |
| *w*<sub>Successful</sub> | 2.74 | [1.80, 3.73] | 0.88 | 0.26 | 0.57 | 1.00 |
| *w*<sub>Unsuccessful</sub> | -2.00 | [-2.64, -1.38] | 0.53 | 0.51 | 1.34 | 1.00 |
| *w*<sub>Social</sub> | 0.73 | [0.52, 0.94] | 0.16 | 0.24 | 0.18 | 1.00 |
| *w*<sub>Prior</sub> | 3.36 | [1.94, 4.86] | 0.82 | 0.49 | 1.01 | 1.00 |
| $w_{\mathrm{Successful}\times T_A}$ | -1.21 | [-2.19, -0.01] | 0.81 | 0.19 | 1.94 | 1.00 |
| $w_{\mathrm{Unsuccessful}\times T_A}$ | 1.71 | [1.11, 2.30] | 0.45 | 0.19 | 1.64 | 1.00 |
| $w_{\mathrm{Social}\times T_A}$ | 0.27 | [-0.10, 0.65] | 0.28 | 0.33 | 0.57 | 1.00 |
| $w_{\mathrm{Prior}\times T_A}$ | -3.23 | [-5.61, -0.81] | 0.82 | 0.66 | 1.81 | 1.00 |
| *w*<sub>Successful×Prior</sub> | -0.29 | [-1.10, 0.58] | 0.36 | 0.34 | 0.76 | 1.00 |
| *w*<sub>Unsuccessful×Prior</sub> | 0.21 | [-0.37, 0.80] | 0.30 | 0.53 | 1.09 | 1.00 |
| *w*<sub>Social×Prior</sub> | -0.17 | [-0.50, 0.15] | 0.20 | 0.23 | 0.30 | 1.00 |
| λ<sub>Social</sub> | 3.07 | [2.89, 3.25] | 0.17 | n.a. | n.a. | 1.00 |
| λ<sub>Personal</sub> | 2.55 | [2.33, 2.81] | 0.23 | n.a. | n.a. | 1.00 |

Note. post. mean = posterior mean; CrI = credible interval; σ refers to the posterior estimated standard deviation of parameter values at the respective hierarchical level (Lake, N = 5; Id, N = 62; Trip, N = 251); $\hat{R}$ measures chain convergence with values close to 1 indicating chain convergence.

**Supplementary Table S6.**
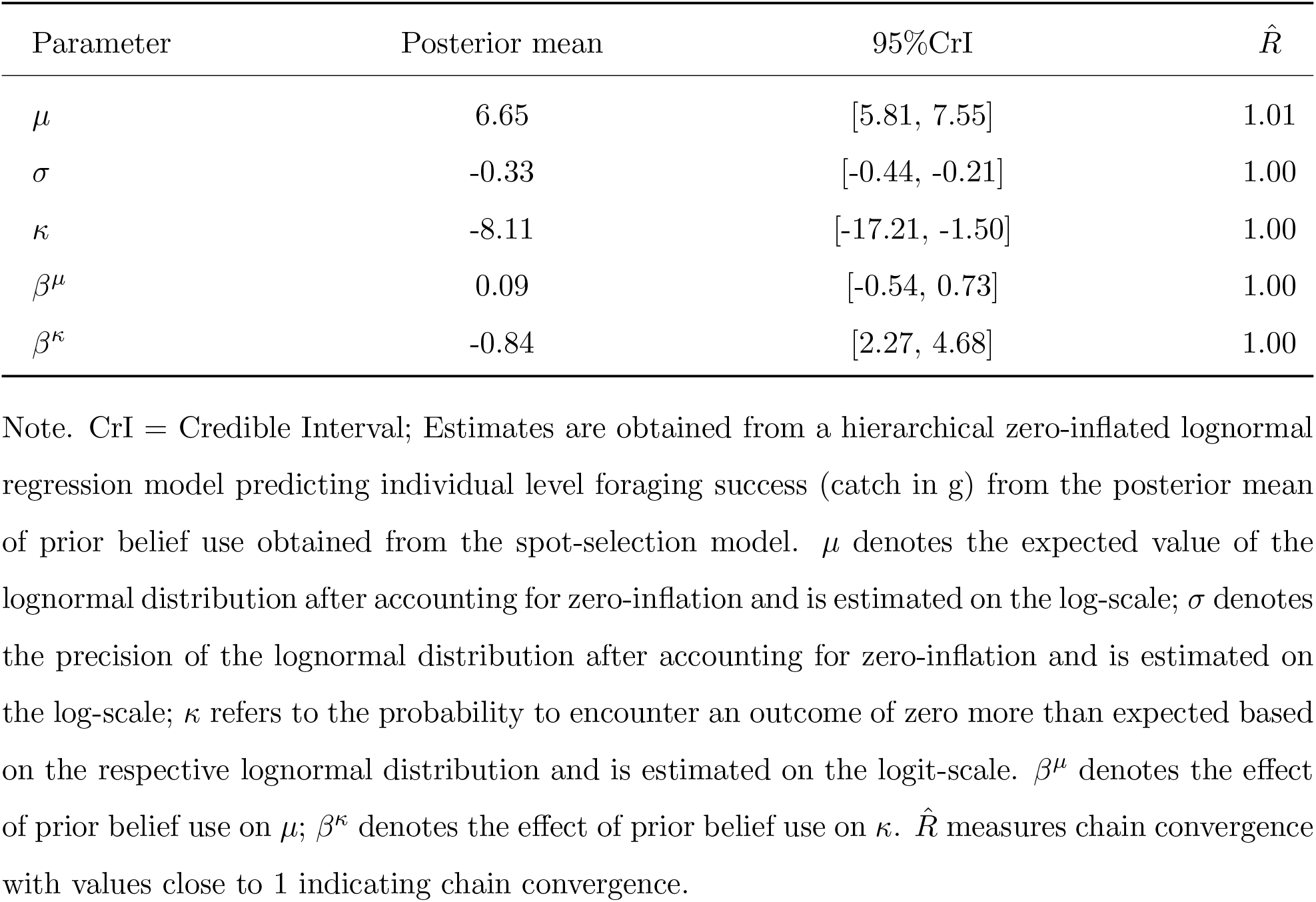
Posterior Estimates Predicting Individual-Level Foraging Success (catch in g) from Prior Belief Use for Spot-Selection.

| Parameter | Posterior mean | 95%CrI | R̂ |
|---|---|---|---|
| μ | 6.65 | [5.81, 7.55] | 1.01 |
| σ | -0.33 | [-0.44, -0.21] | 1.00 |
| κ | -8.11 | [-17.21, -1.50] | 1.00 |
| β<sup>μ</sup> | 0.09 | [-0.54, 0.73] | 1.00 |
| β<sup>κ</sup> | -0.84 | [2.27, 4.68] | 1.00 |

Note. CrI = Credible Interval; Estimates are obtained from a hierarchical zero-inflated lognormal regression model predicting individual level foraging success (catch in g) from the posterior mean of prior belief use obtained from the spot-selection model. μ denotes the expected value of the lognormal distribution after accounting for zero-inflation and is estimated on the log-scale; σ denotes the precision of the lognormal distribution after accounting for zero-inflation and is estimated on the log-scale; κ refers to the probability to encounter an outcome of zero more than expected based on the respective lognormal distribution and is estimated on the logit-scale. β<sup>μ</sup> denotes the effect of prior belief use on μ; β<sup>κ</sup> denotes the effect of prior belief use on κ. R̂ measures chain convergence with values close to 1 indicating chain convergence.

**Supplementary Table S7.**
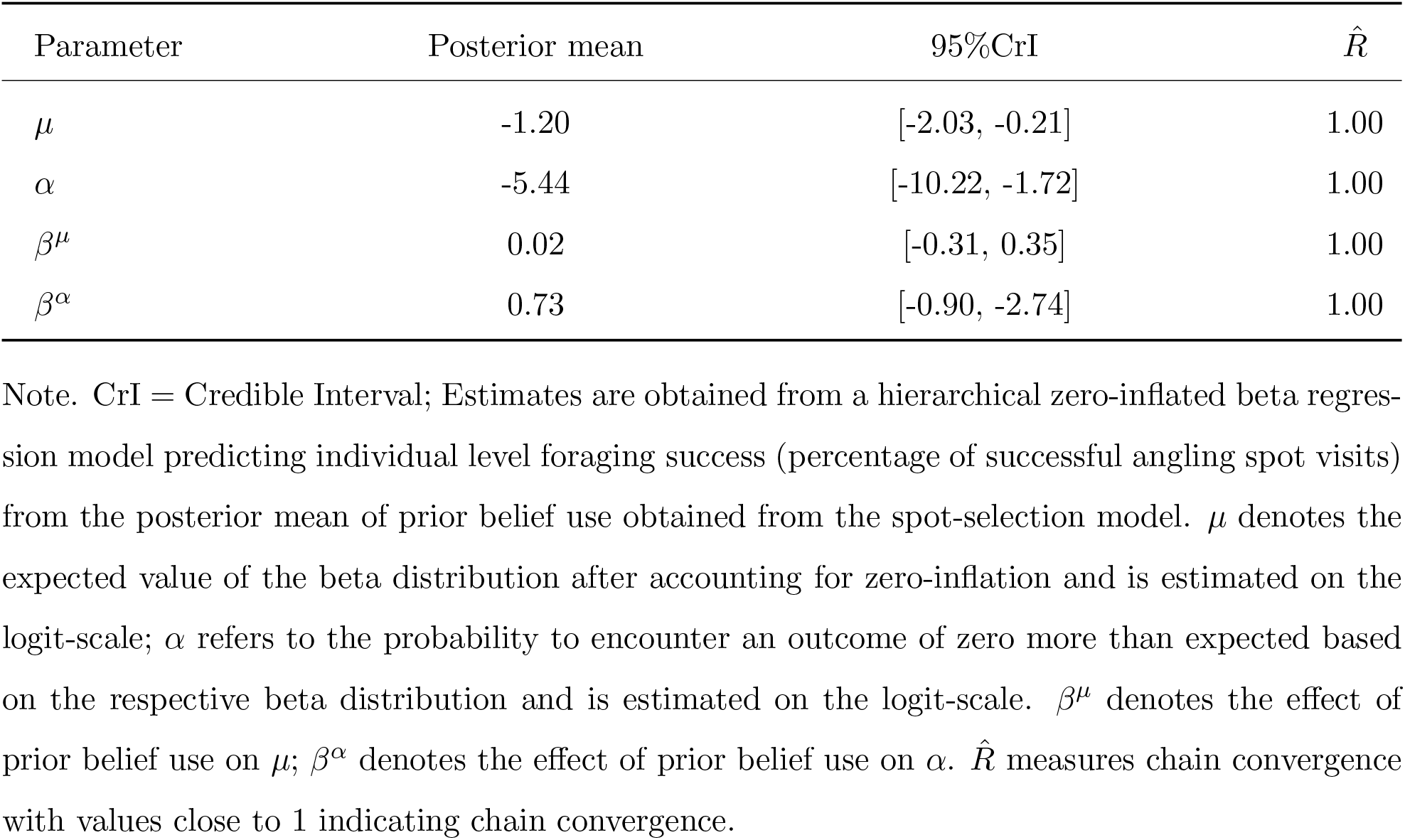
Posterior Estimates Predicting Individual-Level Foraging Success (% Successful Spots) From Prior Belief Use for Spot-Selection.

| Parameter | Posterior mean | 95%CrI | R̂ |
|---|---|---|---|
| μ | -1.20 | [-2.03, -0.21] | 1.00 |
| α | -5.44 | [-10.22, -1.72] | 1.00 |
| β<sup>μ</sup> | 0.02 | [-0.31, 0.35] | 1.00 |
| β<sup>α</sup> | 0.73 | [-0.90, -2.74] | 1.00 |

Note. CrI = Credible Interval; Estimates are obtained from a hierarchical zero-inflated beta regres-sion model predicting individual level foraging success (percentage of successful angling spot visits) from the posterior mean of prior belief use obtained from the spot-selection model. μ denotes the expected value of the beta distribution after accounting for zero-inflation and is estimated on the logit-scale; α refers to the probability to encounter an outcome of zero more than expected based on the respective beta distribution and is estimated on the logit-scale. β<sup>μ</sup> denotes the effect of prior belief use on μ; β<sup>α</sup> denotes the effect of prior belief use on α. R̂ measures chain convergence with values close to 1 indicating chain convergence.

**Supplementary Table S8.**
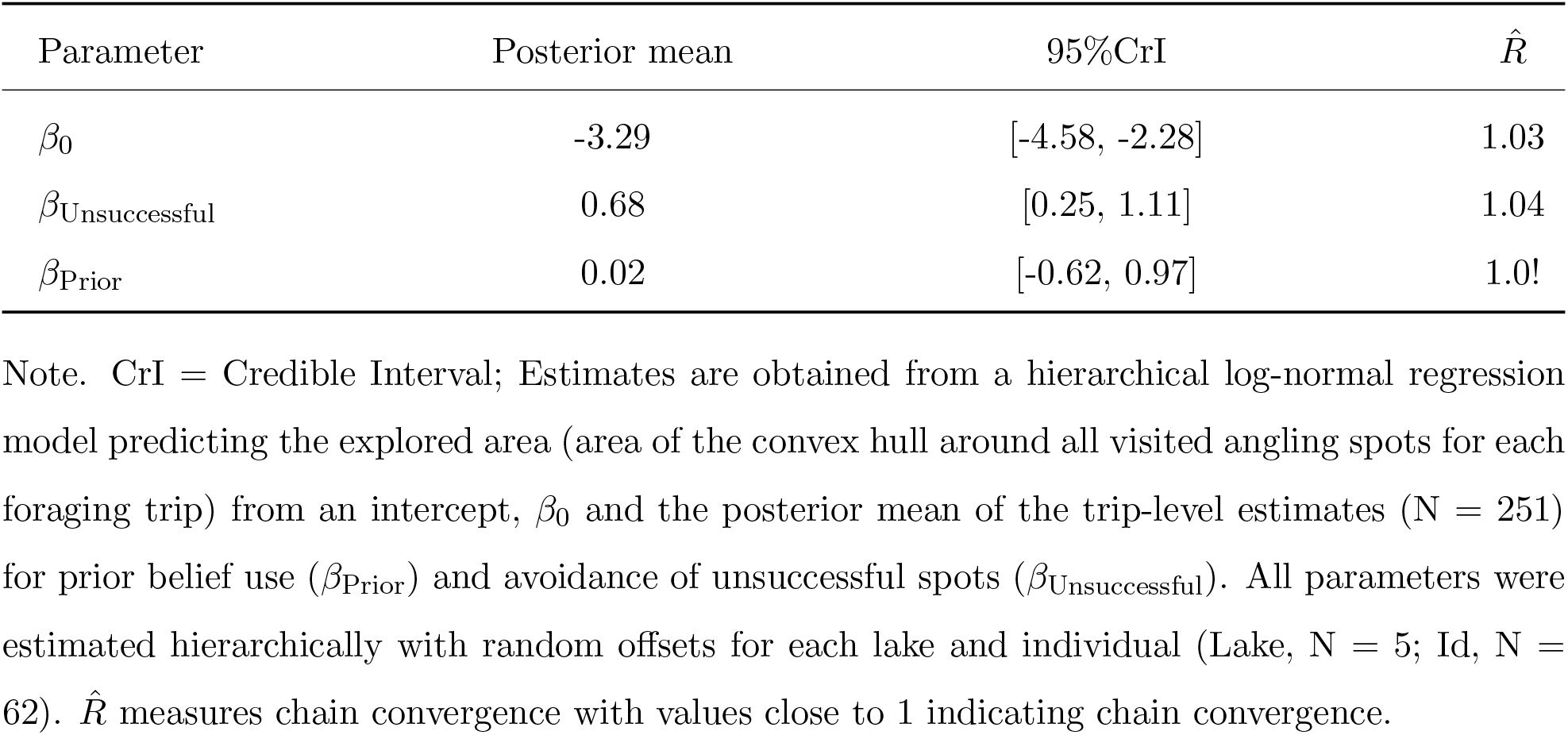
Posterior Estimates Predicting Explored Area from Prior Belief Use and Avoidance of Unsuccessful Spots for Spot-Selection.

| Parameter | Posterior mean | 95%CrI | $\hat{R}$ |
|---|---|---|---|
| β<sub>0</sub> | -3.29 | [-4.58, -2.28] | 1.03 |
| β<sub>Unsuccessful</sub> | 0.68 | [0.25, 1.11] | 1.04 |
| β<sub>Prior</sub> | 0.02 | [-0.62, 0.97] | 1.0! |

Note. CrI = Credible Interval; Estimates are obtained from a hierarchical log-normal regression model predicting the explored area (area of the convex hull around all visited angling spots for each foraging trip) from an intercept, β<sub>0</sub> and the posterior mean of the trip-level estimates (N = 251) for prior belief use (β<sub>Prior</sub>) and avoidance of unsuccessful spots (β<sub>Unsuccessful</sub>). All parameters were estimated hierarchically with random offsets for each lake and individual (Lake, N = 5; Id, N = 62). $\hat{R}$ measures chain convergence with values close to 1 indicating chain convergence.

**Supplementary Table S9.**
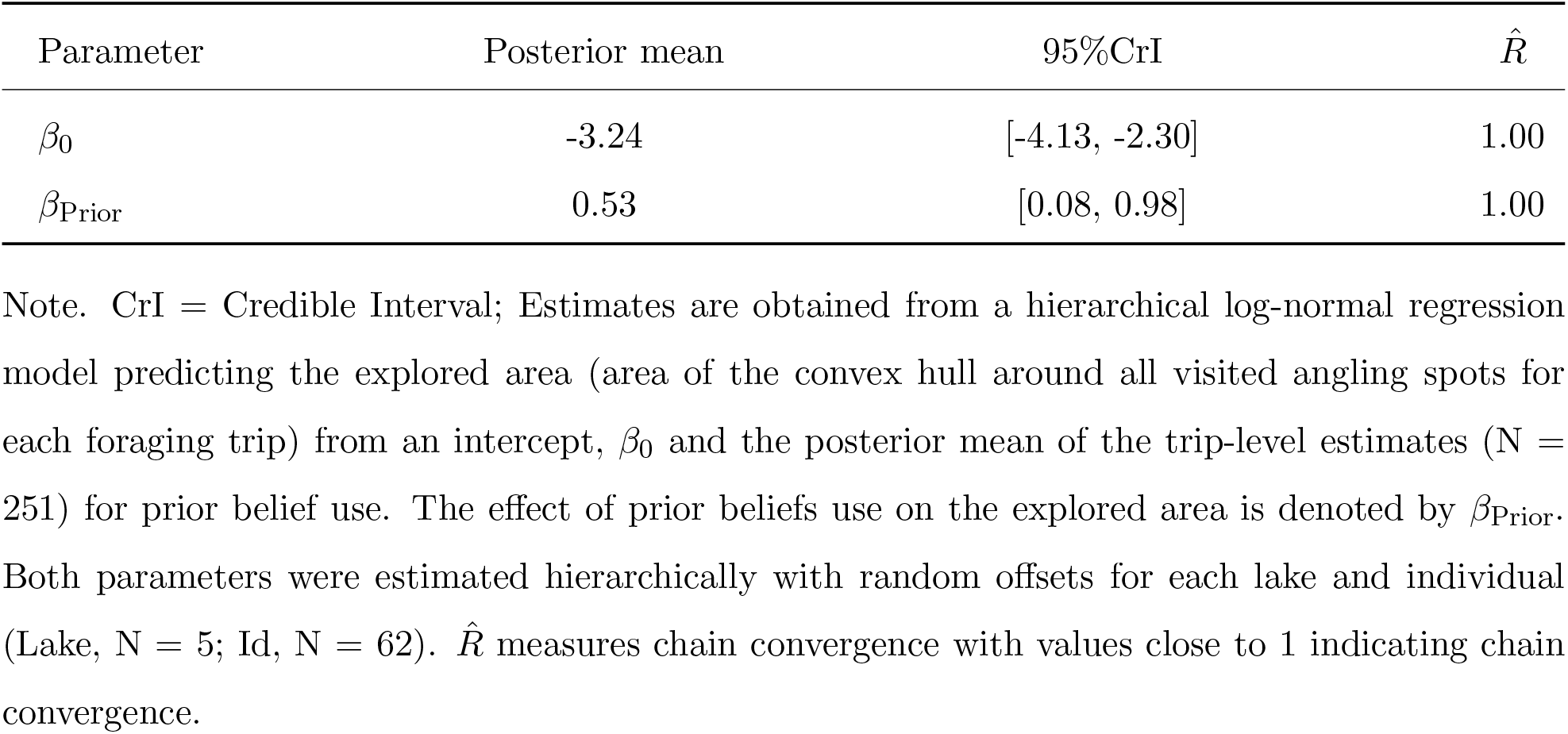
Posterior Estimates Predicting Explored Area from Prior Beliefs.

| Parameter | Posterior mean | 95%CrI | R̂ |
|---|---|---|---|
| β<sub>0</sub> | -3.24 | [-4.13, -2.30] | 1.00 |
| β<sub>Prior</sub> | 0.53 | [0.08, 0.98] | 1.00 |

Note. CrI = Credible Interval; Estimates are obtained from a hierarchical log-normal regression model predicting the explored area (area of the convex hull around all visited angling spots for each foraging trip) from an intercept, β<sub>0</sub> and the posterior mean of the trip-level estimates (N = 251) for prior belief use. The effect of prior beliefs use on the explored area is denoted by β<sub>Prior</sub>. Both parameters were estimated hierarchically with random offsets for each lake and individual (Lake, N = 5; Id, N = 62). R̂ measures chain convergence with values close to 1 indicating chain convergence.

**Supplementary Table S10.**
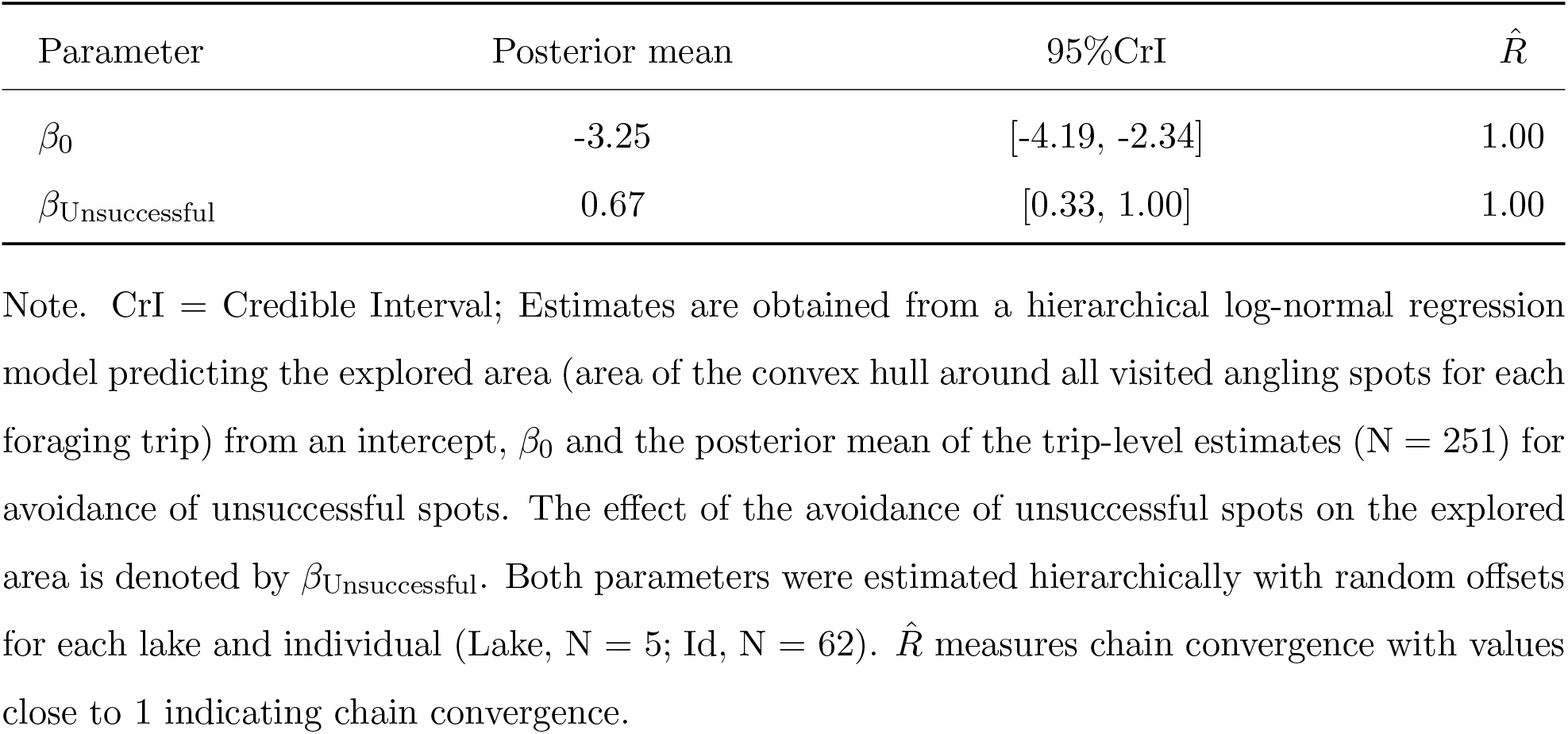
Posterior Estimates Predicting Explored Area from Loss Avoidance.

| Parameter | Posterior mean | 95%CrI | R̂ |
|---|---|---|---|
| β<sub>0</sub> | -3.25 | [-4.19, -2.34] | 1.00 |
| β<sub>Unsuccessful</sub> | 0.67 | [0.33, 1.00] | 1.00 |

Note. CrI = Credible Interval; Estimates are obtained from a hierarchical log-normal regression model predicting the explored area (area of the convex hull around all visited angling spots for each foraging trip) from an intercept, β<sub>0</sub> and the posterior mean of the trip-level estimates (N = 251) for avoidance of unsuccessful spots. The effect of the avoidance of unsuccessful spots on the explored area is denoted by β<sub>Unsuccessful</sub>. Both parameters were estimated hierarchically with random offsets for each lake and individual (Lake, N = 5; Id, N = 62). R̂ measures chain convergence with values close to 1 indicating chain convergence.

**Supplementary Table S11.** Unstandardized Posterior Estimates of the Spot-Leaving Model.

| Parameter | Post. mean | 95%CrI | *σ*<sub>Lake</sub> | *σ*<sub>Trip</sub> | *σ*<sub>ID</sub> | $\hat{R}$ |
|---|---|---|---|---|---|---|
| *w*<sub>0</sub> | -3.64 | [-4.03, -3.21] | 0.39 | 0.41 | 0.35 | 1.00 |
| *w*<sub>Fish</sub> | -2.10 | [-2.81, -1.36] | 0.71 | 0.32 | 0.85 | 1.00 |
| *w*<sub>Time</sub> | 0.13 | [0.05, 0.21] | 0.08 | 0.06 | 0.07 | 1.00 |
| *w*<sub>Successful</sub> | 0.08 | [-0.14, 0.28] | 0.18 | 0.11 | 0.26 | 1.00 |
| *w*<sub>Unsuccessful</sub> | 0.01 | [-0.18, 0.20] | 0.18 | 0.05 | 0.07 | 1.00 |
| *w*<sub>Prior</sub> | 0.03 | [-0.11, 0.18] | 0.11 | 0.08 | 0.23 | 1.00 |
| *w*<sub>Social</sub> | -0.37 | [-0.64, -0.13] | 0.22 | 0.13 | 0.33 | 1.00 |

Note. post. mean = posterior mean; CrI = credible interval; *σ* refers to the posterior estimated standard deviation of parameter values at the respective hierarchical level (Lake, N = 5; Id, N = 62; Trip, N = 251); $\hat{R}$ measures chain convergence with values close to 1 indicating chain convergence. Feature weights are referred to as *w* with the subscript denoting the respective feature: *w*<sub>0</sub> denotes the intercept, *w*<sub>Time</sub> denotes the effect of time without fish catches, *w*<sub>Fish</sub> denotes the effect of finding the first fish, *w*<sub>Successful</sub>, *w*<sub>Unsuccessful</sub>, *w*<sub>Social</sub> and *w*<sub>Prior</sub> denote the effects of the four spatial features on the probability to leave (on the logit scale). $\hat{R}$ measures chain convergence with values close to 1 indicating chain convergence.

**Supplementary Table S12.**
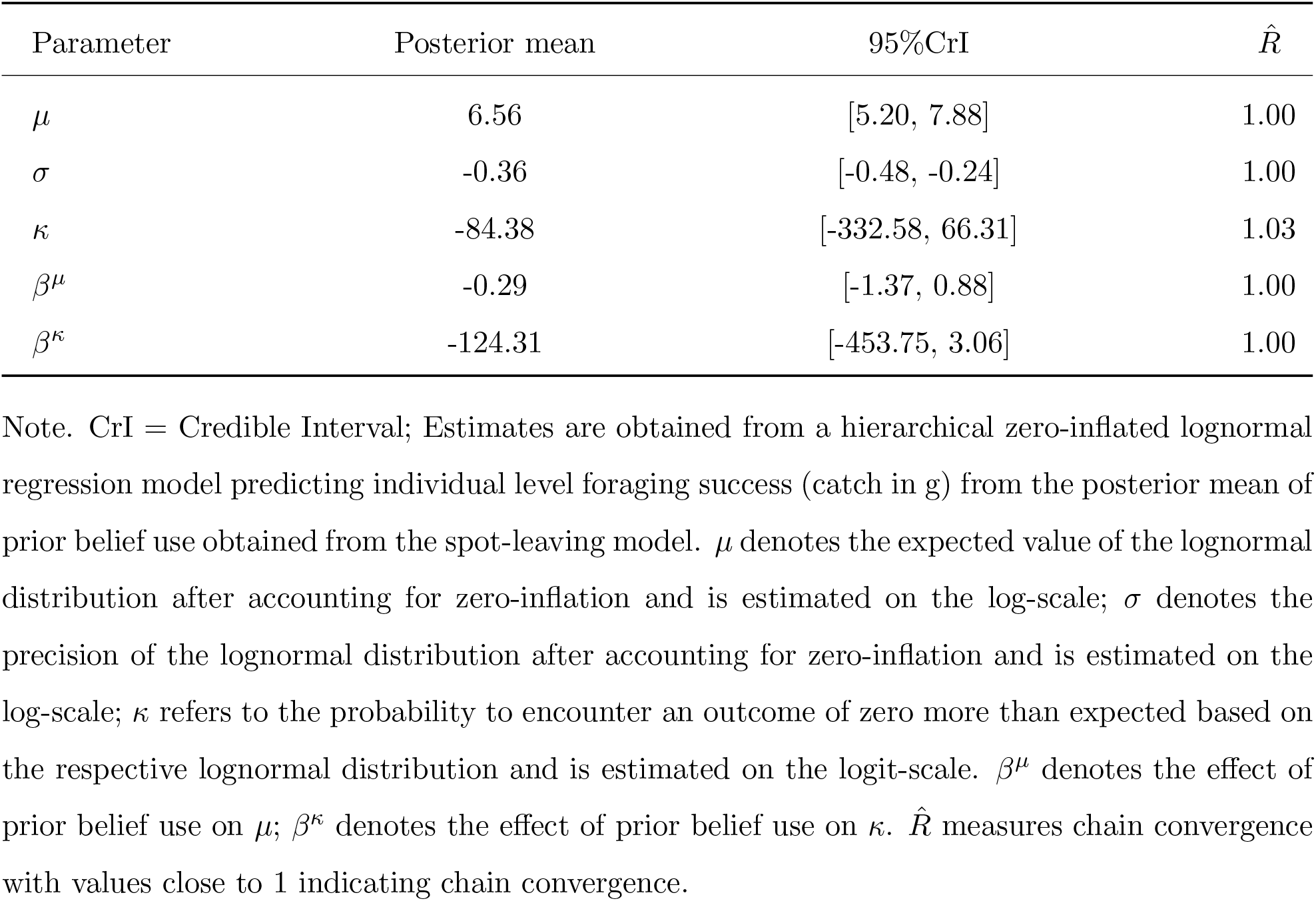
Posterior Estimates Predicting Individual-Level Foraging Success (catch in g) From Prior Belief Use for Spot-Leaving.

| Parameter | Posterior mean | 95%CrI | R̂ |
|---|---|---|---|
| μ | 6.56 | [5.20, 7.88] | 1.00 |
| σ | -0.36 | [-0.48, -0.24] | 1.00 |
| κ | -84.38 | [-332.58, 66.31] | 1.03 |
| β<sup>μ</sup> | -0.29 | [-1.37, 0.88] | 1.00 |
| β<sup>κ</sup> | -124.31 | [-453.75, 3.06] | 1.00 |

Note. CrI = Credible Interval; Estimates are obtained from a hierarchical zero-inflated lognormal regression model predicting individual level foraging success (catch in g) from the posterior mean of prior belief use obtained from the spot-leaving model. μ denotes the expected value of the lognormal distribution after accounting for zero-inflation and is estimated on the log-scale; σ denotes the precision of the lognormal distribution after accounting for zero-inflation and is estimated on the log-scale; κ refers to the probability to encounter an outcome of zero more than expected based on the respective lognormal distribution and is estimated on the logit-scale. β<sup>μ</sup> denotes the effect of prior belief use on μ; β<sup>κ</sup> denotes the effect of prior belief use on κ. R̂ measures chain convergence with values close to 1 indicating chain convergence.

**Supplementary Table S13.**
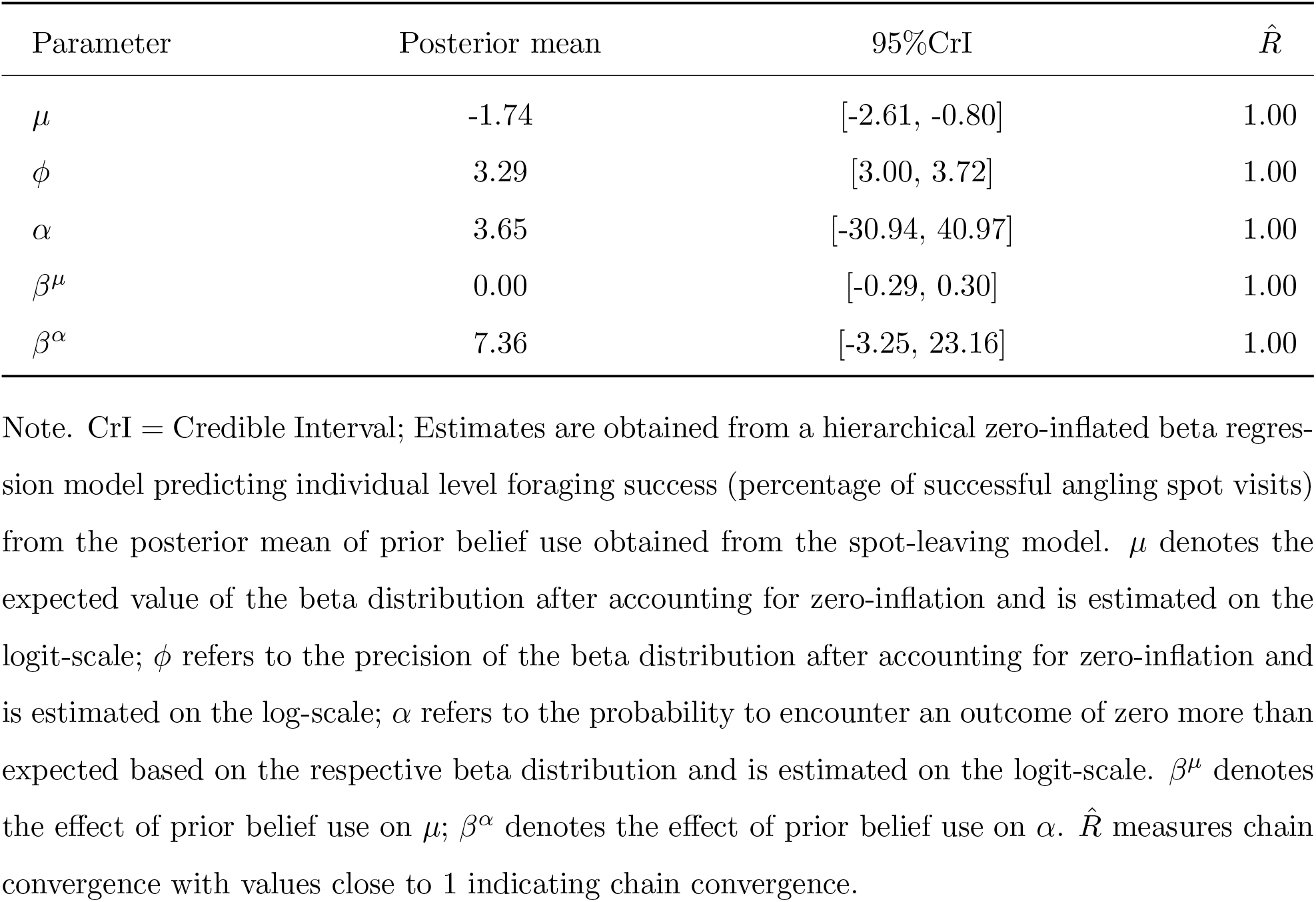
Posterior Estimates Predicting Individual-Level Foraging Success (% Successful Spots) From Prior Belief Use for Spot-Leaving.

| Parameter | Posterior mean | 95%CrI | R̂ |
|---|---|---|---|
| μ | -1.74 | [-2.61, -0.80] | 1.00 |
| ϕ | 3.29 | [3.00, 3.72] | 1.00 |
| α | 3.65 | [-30.94, 40.97] | 1.00 |
| β<sup>μ</sup> | 0.00 | [-0.29, 0.30] | 1.00 |
| β<sup>α</sup> | 7.36 | [-3.25, 23.16] | 1.00 |

Note. CrI = Credible Interval; Estimates are obtained from a hierarchical zero-inflated beta regres-sion model predicting individual level foraging success (percentage of successful angling spot visits) from the posterior mean of prior belief use obtained from the spot-leaving model. μ denotes the expected value of the beta distribution after accounting for zero-inflation and is estimated on the logit-scale; ϕ refers to the precision of the beta distribution after accounting for zero-inflation and is estimated on the log-scale; α refers to the probability to encounter an outcome of zero more than expected based on the respective beta distribution and is estimated on the logit-scale. β<sup>μ</sup> denotes the effect of prior belief use on μ; β<sup>α</sup> denotes the effect of prior belief use on α. R̂ measures chain convergence with values close to 1 indicating chain convergence.

**Supplementary Table S14.**
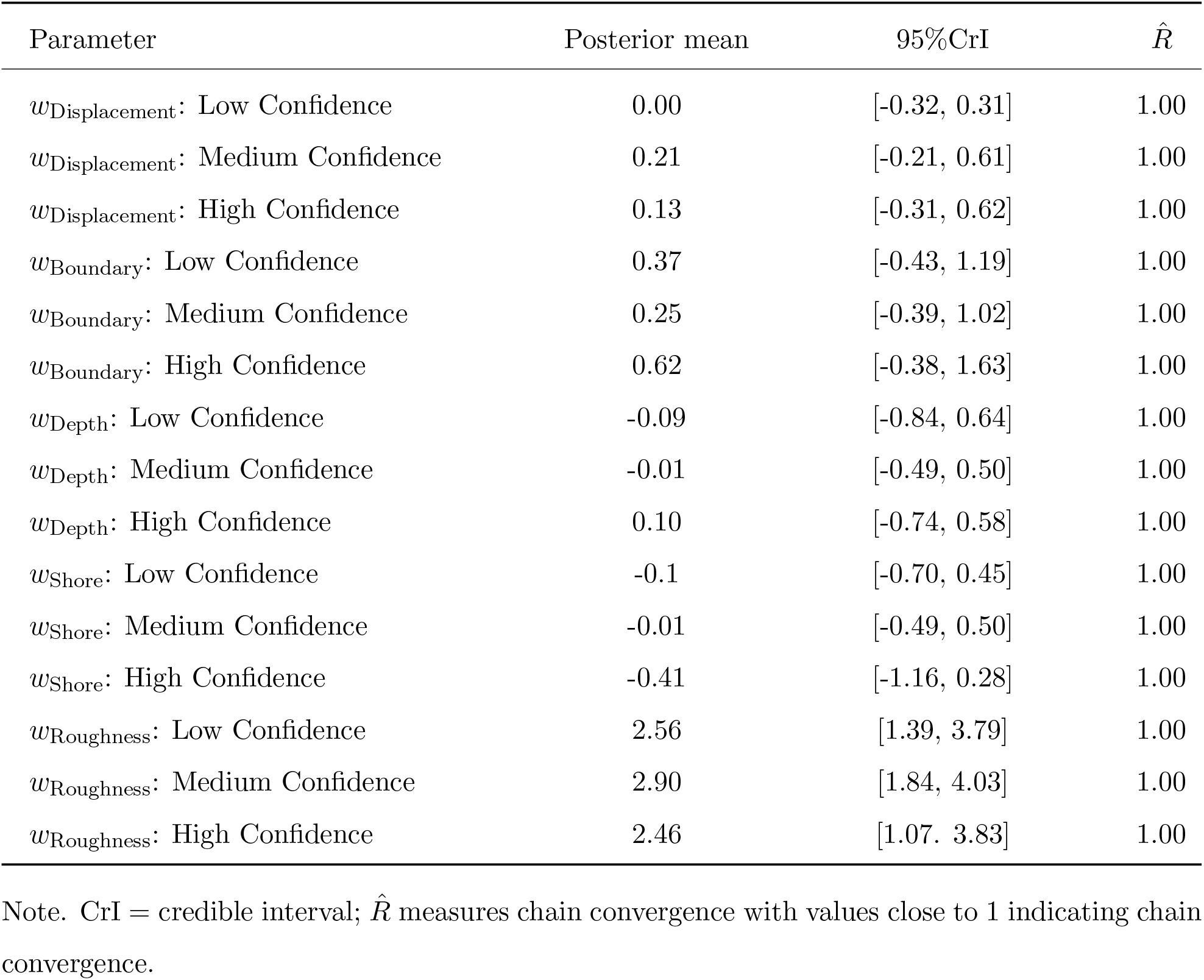
Standardized Posterior Estimates and Convergence Diagnostics of the Confidence Effects of the Area-Nomination Model.

| Parameter | Posterior mean | 95%CrI | *R̂* |
|---|---|---|---|
| *w*<sub>Displacement</sub>: Low Confidence | 0.00 | [-0.32, 0.31] | 1.00 |
| *w*<sub>Displacement</sub>: Medium Confidence | 0.21 | [-0.21, 0.61] | 1.00 |
| *w*<sub>Displacement</sub>: High Confidence | 0.13 | [-0.31, 0.62] | 1.00 |
| *w*<sub>Boundary</sub>: Low Confidence | 0.37 | [-0.43, 1.19] | 1.00 |
| *w*<sub>Boundary</sub>: Medium Confidence | 0.25 | [-0.39, 1.02] | 1.00 |
| *w*<sub>Boundary</sub>: High Confidence | 0.62 | [-0.38, 1.63] | 1.00 |
| *w*<sub>Depth</sub>: Low Confidence | -0.09 | [-0.84, 0.64] | 1.00 |
| *w*<sub>Depth</sub>: Medium Confidence | -0.01 | [-0.49, 0.50] | 1.00 |
| *w*<sub>Depth</sub>: High Confidence | 0.10 | [-0.74, 0.58] | 1.00 |
| *w*<sub>Shore</sub>: Low Confidence | -0.1 | [-0.70, 0.45] | 1.00 |
| *w*<sub>Shore</sub>: Medium Confidence | -0.01 | [-0.49, 0.50] | 1.00 |
| *w*<sub>Shore</sub>: High Confidence | -0.41 | [-1.16, 0.28] | 1.00 |
| *w*<sub>Roughness</sub>: Low Confidence | 2.56 | [1.39, 3.79] | 1.00 |
| *w*<sub>Roughness</sub>: Medium Confidence | 2.90 | [1.84, 4.03] | 1.00 |
| *w*<sub>Roughness</sub>: High Confidence | 2.46 | [1.07. 3.83] | 1.00 |

Note. CrI = credible interval; *R̂* measures chain convergence with values close to 1 indicating chain convergence.

**Supplementary Table S15.**
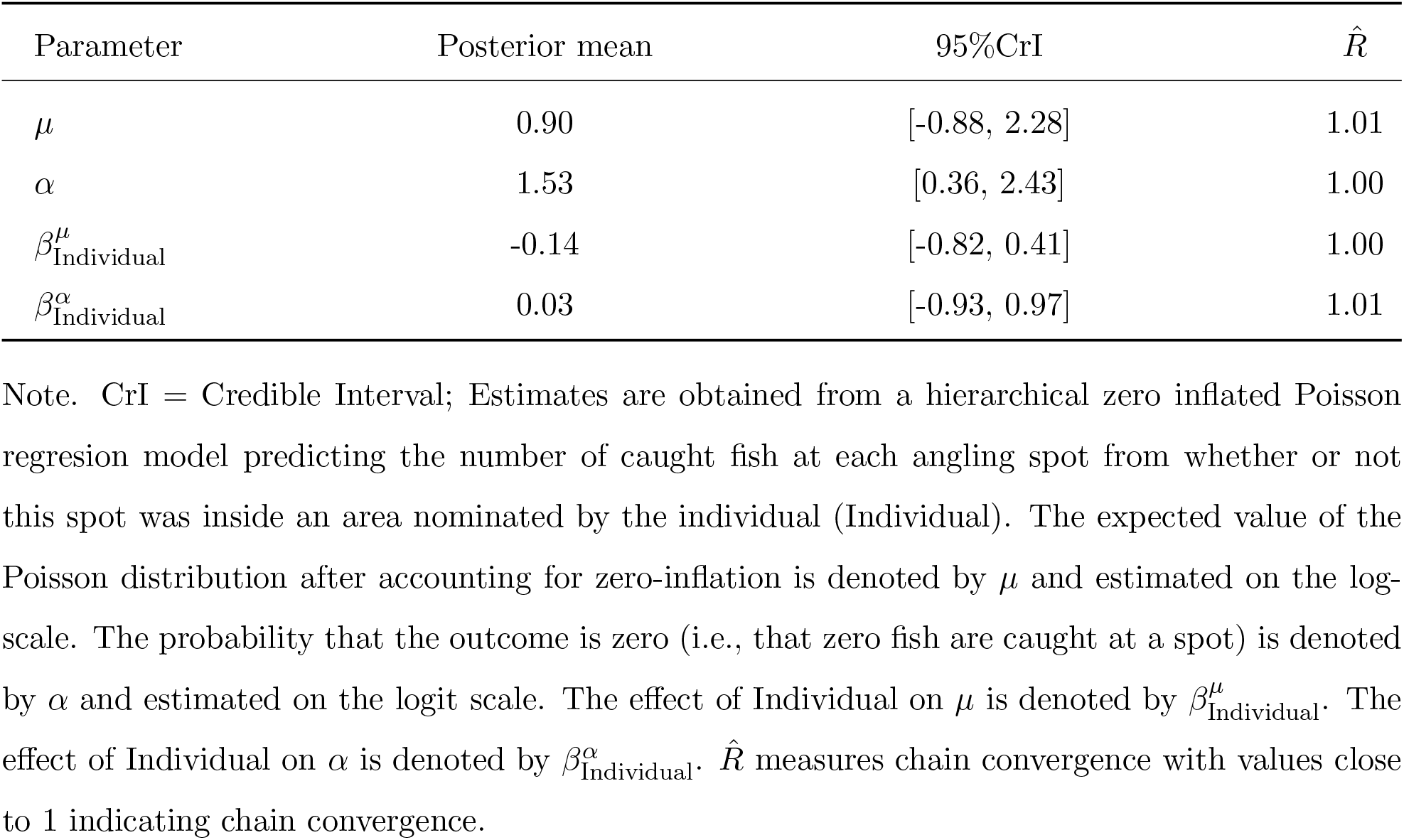
Posterior Estimates Predicting Foraging Success at each Spot from Individually Nominated Areas.

| Parameter | Posterior mean | 95%CrI | R̂ |
|---|---|---|---|
| μ | 0.90 | [-0.88, 2.28] | 1.01 |
| α | 1.53 | [0.36, 2.43] | 1.00 |
| β<sup>μ</sup><sub>Individual</sub> | -0.14 | [-0.82, 0.41] | 1.00 |
| β<sup>α</sup><sub>Individual</sub> | 0.03 | [-0.93, 0.97] | 1.01 |

Note. CrI = Credible Interval; Estimates are obtained from a hierarchical zero inflated Poisson regresion model predicting the number of caught fish at each angling spot from whether or not this spot was inside an area nominated by the individual (Individual). The expected value of the Poisson distribution after accounting for zero-inflation is denoted by μ and estimated on the log-scale. The probability that the outcome is zero (i.e., that zero fish are caught at a spot) is denoted by α and estimated on the logit scale. The effect of Individual on μ is denoted by β<sup>μ</sup><sub>Individual</sub>. The effect of Individual on α is denoted by β<sup>α</sup><sub>Individual</sub>. R̂ measures chain convergence with values close to 1 indicating chain convergence.

**Supplementary Table S16.**
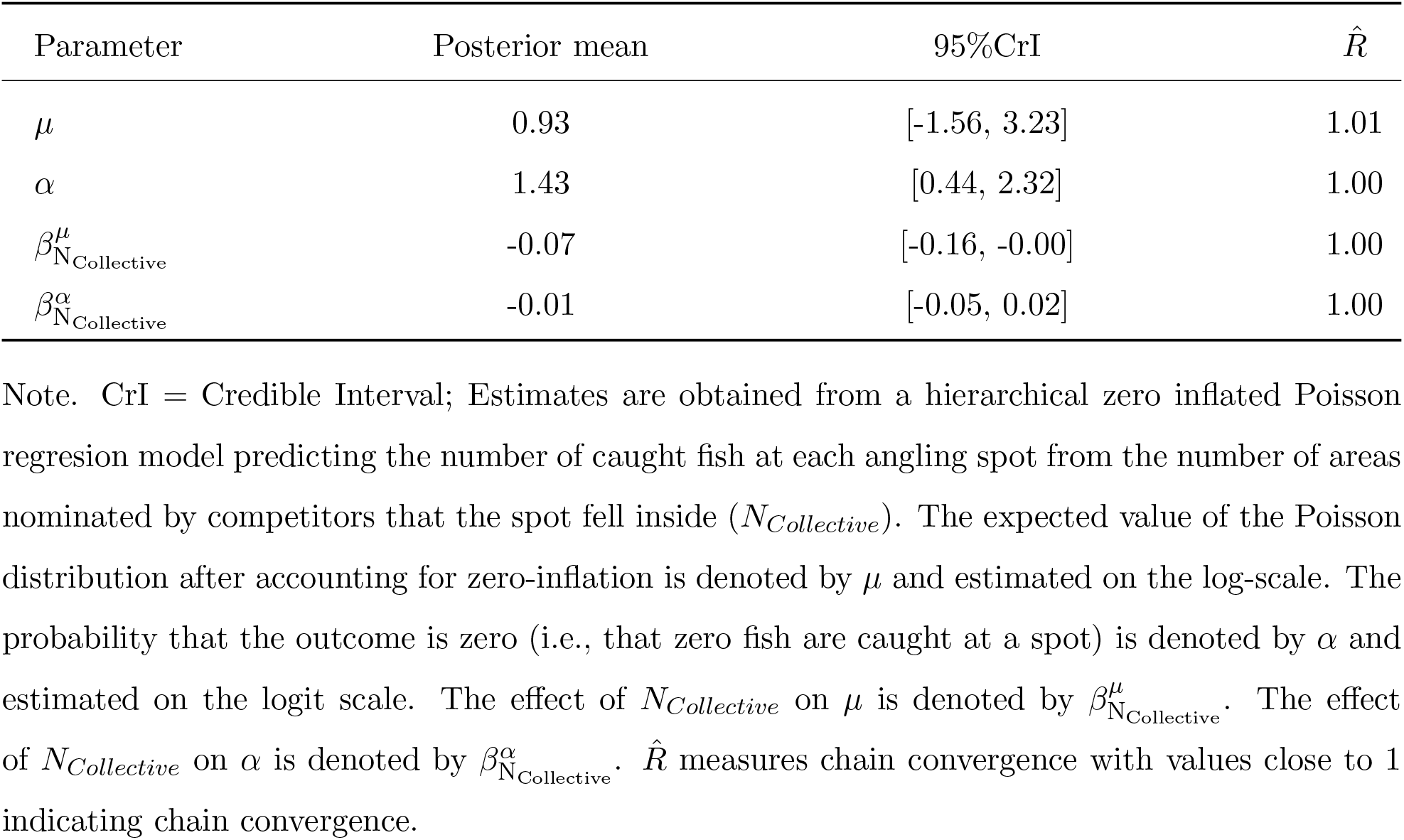
Posterior Estimates Predicting Foraging Success at each Spot from Collectively Nominated Areas.

| Parameter | Posterior mean | 95%CrI | R̂ |
|---|---|---|---|
| μ | 0.93 | [-1.56, 3.23] | 1.01 |
| α | 1.43 | [0.44, 2.32] | 1.00 |
| β<sup>μ</sup><sub>N<sub>Collective</sub></sub> | -0.07 | [-0.16, -0.00] | 1.00 |
| β<sup>α</sup><sub>N<sub>Collective</sub></sub> | -0.01 | [-0.05, 0.02] | 1.00 |

Note. CrI = Credible Interval; Estimates are obtained from a hierarchical zero inflated Poisson regresion model predicting the number of caught fish at each angling spot from the number of areas nominated by competitors that the spot fell inside (*N<sub>Collective</sub>*). The expected value of the Poisson distribution after accounting for zero-inflation is denoted by μ and estimated on the log-scale. The probability that the outcome is zero (i.e., that zero fish are caught at a spot) is denoted by α and estimated on the logit scale. The effect of *N<sub>Collective</sub>* on μ is denoted by β<sup>μ</sup><sub>N<sub>Collective</sub></sub>. The effect of *N<sub>Collective</sub>* on α is denoted by β<sup>α</sup><sub>N<sub>Collective</sub></sub>. R̂ measures chain convergence with values close to 1 indicating chain convergence.

**Supplementary Table S17.** Unstandardized Posterior Estimates and Convergence Diagnostics of Prior Belief Use Depending on Confidence Levels.

| Parameter | Post. mean | 95%CrI | σ<sub>Lake</sub> | σ<sub>Id</sub> | σ<sub>Trip</sub> | $\hat{R}$ |
|---|---|---|---|---|---|---|
| *w*<sub>Prior</sub> High Confidence | 1.24 | [1.35, 2.63] | 0.70 | 0.39 | 1.43 | 1.00 |
| *w*<sub>Prior</sub> Medium Confidence | 1.00 | [0.01, 2.22] | 0.60 | 0.52 | 0.46 | 1.00 |
| *w*<sub>Prior</sub> Low Confidence | 1.94 | [0.85, 3.05] | 0.64 | 0.43 | 0.42 | 1.00 |

Note. Post. mean = posterior mean; CrI = credible interval; σ refers to the posterior estimated standard deviation of parameter values at the respective hierarchical level (Lake, N = 5; Id, N = 62; Trip, N = 251); *w*<sub>Prior</sub> refers to the weight of the inverse distance to the closest nominated area on spot-selection decisions. $\hat{R}$ measures chain convergence with values close to 1 indicating chain convergence.

**Supplementary Table S18.**
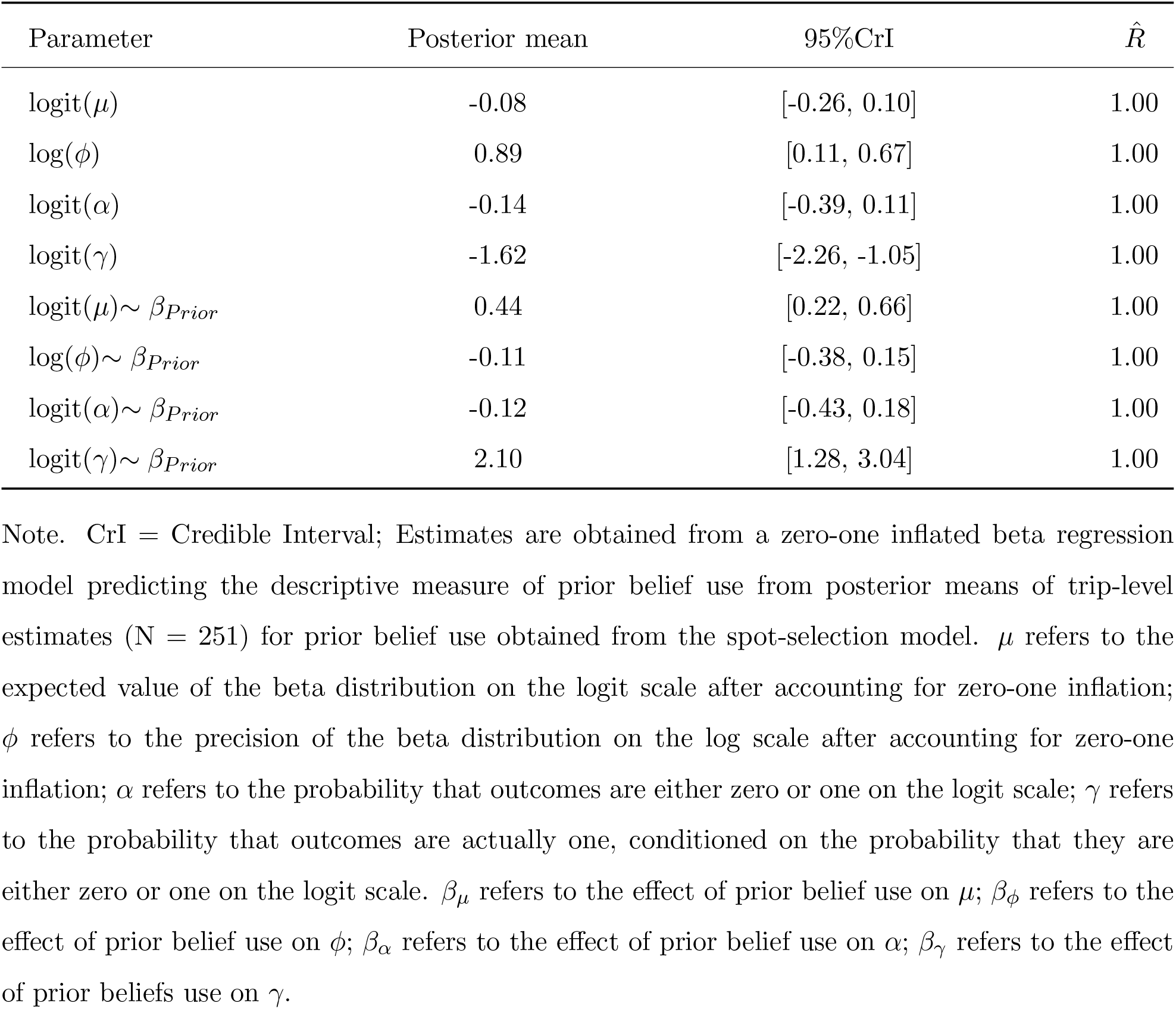
Posterior Estimates Predicting Descriptive Prior Belief Use from Model Estimates for Prior Belief Use.

| Parameter | Posterior mean | 95%CrI | $\hat{R}$ |
|---|---|---|---|
| logit(μ) | -0.08 | [-0.26, 0.10] | 1.00 |
| log(ϕ) | 0.89 | [0.11, 0.67] | 1.00 |
| logit(α) | -0.14 | [-0.39, 0.11] | 1.00 |
| logit(γ) | -1.62 | [-2.26, -1.05] | 1.00 |
| logit(μ)∼ β<sub>Prior</sub> | 0.44 | [0.22, 0.66] | 1.00 |
| log(ϕ)∼ β<sub>Prior</sub> | -0.11 | [-0.38, 0.15] | 1.00 |
| logit(α)∼ β<sub>Prior</sub> | -0.12 | [-0.43, 0.18] | 1.00 |
| logit(γ)∼ β<sub>Prior</sub> | 2.10 | [1.28, 3.04] | 1.00 |

Note. CrI = Credible Interval; Estimates are obtained from a zero-one inflated beta regression model predicting the descriptive measure of prior belief use from posterior means of trip-level estimates (N = 251) for prior belief use obtained from the spot-selection model. μ refers to the expected value of the beta distribution on the logit scale after accounting for zero-one inflation; ϕ refers to the precision of the beta distribution on the log scale after accounting for zero-one inflation; α refers to the probability that outcomes are either zero or one on the logit scale; γ refers to the probability that outcomes are actually one, conditioned on the probability that they are either zero or one on the logit scale. β<sub>μ</sub> refers to the effect of prior belief use on μ; β<sub>ϕ</sub> refers to the effect of prior belief use on ϕ; β<sub>α</sub> refers to the effect of prior belief use on α; β<sub>γ</sub> refers to the effect of prior beliefs use on γ.

**Supplementary Table S19.**
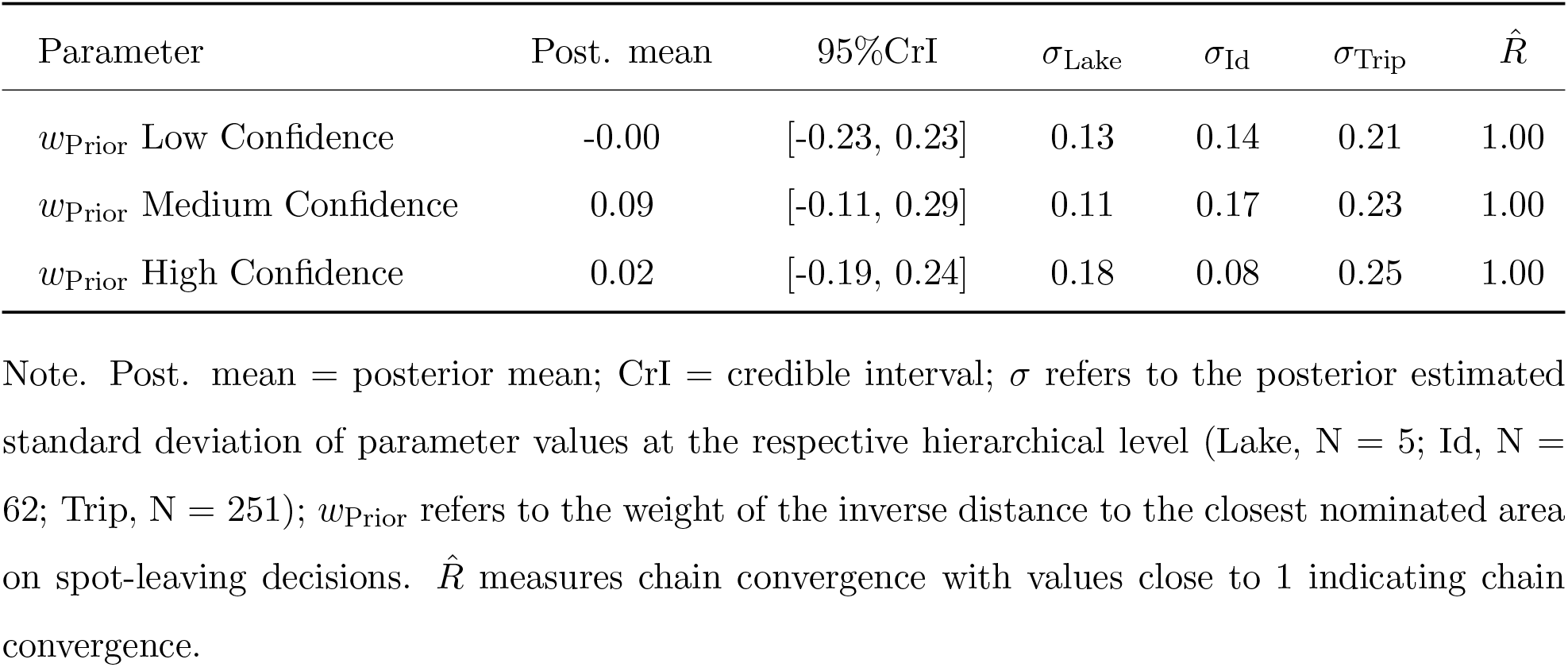
Unstandardized Posterior Estimates of Prior Belief use for Spot-Leaving Decisions Depending on Confidence Levels.

| Parameter | Post. mean | 95%CrI | σ<sub>Lake</sub> | σ<sub>Id</sub> | σ<sub>Trip</sub> | $\hat{R}$ |
|---|---|---|---|---|---|---|
| *w*<sub>Prior</sub> Low Confidence | -0.00 | [-0.23, 0.23] | 0.13 | 0.14 | 0.21 | 1.00 |
| *w*<sub>Prior</sub> Medium Confidence | 0.09 | [-0.11, 0.29] | 0.11 | 0.17 | 0.23 | 1.00 |
| *w*<sub>Prior</sub> High Confidence | 0.02 | [-0.19, 0.24] | 0.18 | 0.08 | 0.25 | 1.00 |

Note. Post. mean = posterior mean; CrI = credible interval; σ refers to the posterior estimated standard deviation of parameter values at the respective hierarchical level (Lake, N = 5; Id, N = 62; Trip, N = 251); *w*<sub>Prior</sub> refers to the weight of the inverse distance to the closest nominated area on spot-leaving decisions. $\hat{R}$ measures chain convergence with values close to 1 indicating chain convergence.

